# Exploring the role of transposable elements to sex gap in longevity in *Drosophila* species

**DOI:** 10.1101/2025.01.22.634250

**Authors:** Miriam Merenciano, Sonia Janillon, Camille Mermet-Bouvier, Nelly Burlet, Valentina Rodriguez-Rada, Victor Ronget, Patricia Gibert, Jean-Michel Gaillard, Jean-François Lemaitre, Gabriel AB Marais, Matthieu Boulesteix, Marie Fablet, Cristina Vieira

## Abstract

In *Drosophila*, like in many other animal species, females tend to live longer than males, a phenomenon known as sex gap in longevity (SGL). One possible explanation for this phenomenon could be related to the activity of transposable elements (TE), which may be present at higher levels in the heterogametic sex. TE activity is normally repressed by epigenetic mechanisms, but this regulation weakens with age. Sex chromosomes, such as the Y chromosome, are enriched in TEs, and age-related TE activity may therefore be more pronounced in older males than in older females, likely affecting longevity patterns. Using three *Drosophila* species, we show that SGL varies naturally among wild-type populations, reflecting both intra-and inter-species differences. Transcriptomic data revealed increased age-related TE expression in *D. melanogaster* and *D. suzukii* flies, but not in *D. simulans*. Moreover, we observed a higher number of upregulated TE copies in older males compared to older females across all the three species tested. Additionally, we detected an increase in TE-chimeric transcript generation in some aged samples, particularly in *D. suzukii* males. Finally, the replacement of the Y chromosome between strains with different SGL led to a progressive reduction in male lifespan and increased TE transcriptional release over generations, suggesting a Y chromosome important role in male longevity. Our work highlights the importance of investigating the role of TEs to better understand differences in longevity between sexes and across different species.

## INTRODUCTION

In most animal species, males and females display distinct mortality patterns leading to differences in their longevity (Marais, et al. 2018; Lemaître, Ronget and Gaillard 2020). This phenomenon is known as sex gap in longevity (SGL) and several hypotheses have been proposed to explain its evolution across the tree of life (Brooks and Garratt 2017; Marais, et al. 2018; Bronikowski, et al. 2022). Among these hypotheses, the putative role played by sex-specific allocation to sexual competition (sexual selection hypothesis) or the asymmetric inheritance of cytoplasmic genomes (mother’s curse hypothesis) have been extensively studied (Frank and Hurst 1996; Vinogradov 1998; Maklakov, et al. 2008; Camus, et al. 2012; Brooks and Garratt 2017; Marais, et al. 2018; Cayuela, et al. 2023). Sex chromosomes have also been proposed to play an important role in SGL since the heterogametic sex generally has the shortest lifespan (Pipoly, et al. 2015; Carazo, et al. 2016; Marais, et al. 2018; Brown, et al. 2020b; Xirocostas, et al. 2020; Nguyen and Bachtrog 2021; Peona, et al. 2021; Marais and Lemaître 2022; Sultanova, et al. 2023). On the one hand, the hemizygosity of the X (in XX/XY taxa) or Z (in ZW/ZZ taxa) chromosome suggests that the shortened lifespan of the heterogametic sex might be due to the unconditional expression of recessive deleterious alleles on its single X (or Z) chromosome (unguarded X effect). However, if the unguarded X effect contributes to sex differences in lifespan is still under debate (Brengdahl, et al. 2018; Connallon, et al. 2022). Alternatively, the toxic Y effect presents another possible hypothesis underlying SGL, postulating that the elevated transposable element (TE) abundance of this chromosome (or W chromosome) compared to other chromosomes may have detrimental consequences for the heterogametic sex (Brown, et al. 2020b; Nguyen and Bachtrog 2021; Peona, et al. 2021; Warmuth, et al. 2022). Y (or W) chromosomes often display reduced rates of recombination, resulting in the accumulation of repeated sequences (mainly TEs) on these chromosomes (Erlandsson, et al. 2000).

TEs are genomic fragments with the ability to move along the genome, being major components of eukaryotic genomes (Bourque, et al. 2018). Due to their mobile and repetitive nature, they are considered a source of mutations and genetic polymorphisms (Bourque, et al. 2018). As such, TE activity is normally repressed by epigenetic regulation that involves DNA methylation, histone modifications and/or small RNAs (Slotkin and Martienssen 2007; Merenciano, et al. 2024). However, with age, these silencing mechanisms are disrupted causing a general increase of TE activity (Wood and Helfand 2013; Copley and Shorter 2023; Yushkova and Moskalev 2023). The toxic Y effect thus proposes that because of the high abundance of TEs on the Y chromosome, more TEs may become active in old males than in old females, which might contribute to sex-differences in ageing and in the reduced male lifespan. Because TEs on the Y chromosome are mostly truncated and non-functional, the impact of TEs on longevity is likely a global effect of the mobilome rather than being driven by Y-linked insertions specifically.

Age-dependent activation of TEs has been suggested to negatively impact lifespan in *Drosophila* (Driver and McKechnie 1992; Nikitin and Woodruff 1995; Woodruff and Nikitin 1995; Li, et al. 2013; Wood and Helfand 2013; Chen, et al. 2016; Wood, et al. 2016; Brown, et al. 2020b; Fabian, et al. 2021; Nguyen and Bachtrog 2021; Rigal, et al. 2022; Yang, et al. 2022), termites (Elsner, et al. 2018; Post, et al. 2023), nematodes (Sturm, et al. 2023), mice (De Cecco, et al. 2019; Ricci, et al. 2023), and humans (Van Meter, et al. 2014; Bogu, et al. 2019; Teoli, et al. 2024; Tsai, et al. 2024). More specifically, in *Drosophila melanogaster*, it has been suggested that aged males underwent a general loss of H3K9me2 histone modification more rapidly than females, resulting in a more pronounced TE activation during ageing in males (Brown, et al. 2020b). Indeed, in this species, it was described that the Y chromosome, and TEs in general, affect chromatin formation genome-wide, in agreement with the heterochromatin sink model (Gatti and Pimpinelli 1992; Brown, et al. 2020a). In addition, longevity assays performed in flies with aberrant sex chromosome configurations (XO, XXY, and XYY) showed that the number of Y chromosomes could influence organismal survival (Brown, et al. 2020b).

In other studies, these results have been challenged. For instance, the generation of *D. melanogaster* Y chromosomes with different sizes using CRISPR/Cas9 suggested that the size of this chromosome, and thus, the amount of heterochromatin (and TEs), did not contribute to sex-specific differences in longevity (Delanoue, et al. 2023). In *D. pseudoobscura*, variations in Y chromosome size and Y-linked TEs showed that larger Y chromosomes lead to slightly increased TE expression, especially in old males, but these differences do not result in faster ageing (Nguyen, et al. 2022). Therefore, the impact of the Y chromosome on male longevity through TE abundance and increased TE expression is still under debate.

In this work, we studied the SGL natural variation across three *Drosophila* species: *D. melanogaster*, *D. simulans*, and *D. suzukii*. Additionally, we analysed TE expression changes associated with longevity in the same species, with particular emphasis on sex-specific differences. Finally, we replaced the Y chromosome among *D. melanogaster* strains to investigate its impact on SGL and lifespan.

## RESULTS

### SGL varies between populations and species

To study the natural variation in SGL, we performed longevity assays in three *Drosophila* species: *D. melanogaster*, *D. simulans*, and *D. suzukii.* For each species, we analyzed between two and six wild-type populations, totaling 28-132 strains, with 100 flies per strain (Fig. 1 and Supplementary Table 1). SGL was measured as the logarithm of the ratio between male and female median lifespan for each strain, a metric previously used in mammals (Lemaître, Ronget, Tidière, et al. 2020) (see Methods) (Supplementary Table 2).

**Figure 1.**
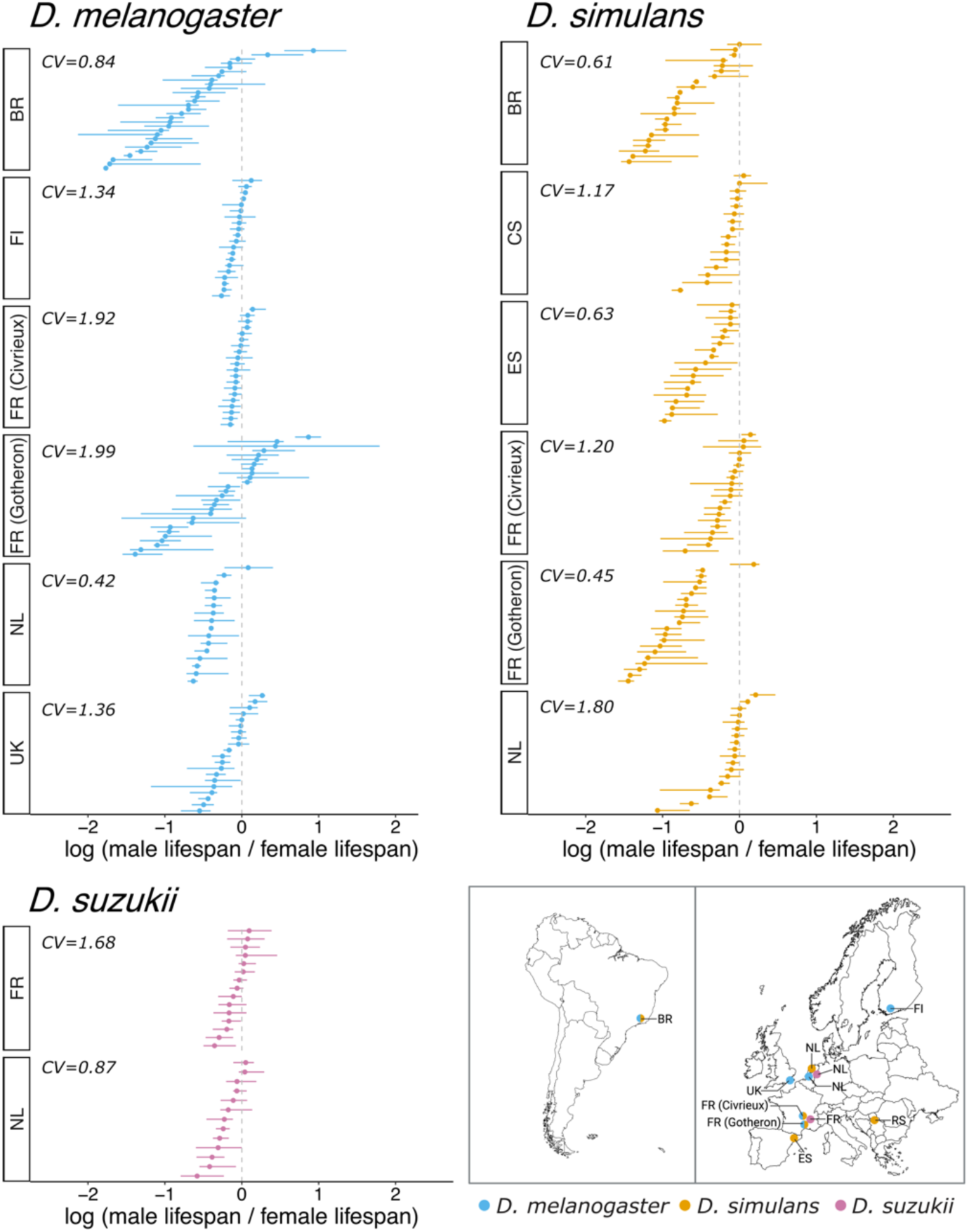
SGL values per strain and geographical distribution of all the populations tested from *D. melanogaster*, *D. simulans*, and *D. suzukii*. Dots show mean SGL values calculated as the logarithm of the ratio between male and female median lifespan and lower and upper whiskers indicate the upper and lower 95% confidence intervals. Coefficients of variation (CV) were calculated for every population. UK: United Kingdom, FR: France, FI: Finland, NL: the Netherlands, BR: Brazil, ES: Spain, RS: Serbia.

In agreement with the literature, the vast majority of strains across all species (82%) exhibited negative SGL values, indicating reduced male longevity (Fig. 1). Specifically, statistically significant negative SGL values were observed in 58% of *D. melanogaster* strains, 65% of *D. simulans* strains, and 35% of *D. suzukii* strains (Fig. 1 and Supplementary Table 2). On average, male lifespan was shorter by 10.5 days in *D. melanogaster*, 12.5 days in *D. simulans*, and nine days in *D. suzukii*.

We then analyzed the variance in the SGL variable across different hierarchical levels (population and species) using a Bayesian approach (see Methods). The analysis revealed that the population of origin contributed significantly to the observed differences in SGL (Supplementary Fig. 1A). Notably, it explained 38.2% (95% CI [16.2, 57.5]) of the total variance suggesting substantial within-species variability attributable to population-level factors (Fig. 1 and Supplementary Fig. 1). In *D. melanogaster*, the coefficient of variation in SGL values ranged from 0.42 to 1.99, while in *D. simulans*, it ranged from 0.45 to 1.80 (Fig. 1). For *D. suzukii*, the coefficients ranged from 0.87 to 1.68, indicating that, although we could analyze only two populations, this species may have the lowest variation in SGL (Fig. 1). The species effect explained 8.6% (95% CI [0.1, 20.8]) of the total variance, also suggesting differences in SGL across species (Fig. 1 and Supplementary Fig. 1B).

In summary, our extensive analysis of SGL showed that males have an overall reduced lifespan compared to females in all three *Drosophila* species tested. Moreover, we found significant within- and between-species variability of SGL values in the wild.

### *D. melanogaster* and *D. suzukii* flies show higher abundance of expressed TE families with age

To study TE expression changes associated with age, we used three *Drosophila* species with varying TE content: *D. melanogaster* (20%), *D. simulans* (12%), and *D. suzukii* (47%) (Mérel, et al. 2020; Mérel, et al. 2021). RNA-seq was performed on young and old individuals from two inbred strains per species, which were previously sequenced with long-read techniques: dmgoth101 and dmgoth63 for *D. melanogaster*, dsgoth31 and dsgoth613 for *D. simulans*, and MT47 and S29 for *D. suzukii* (Mohamed, et al. 2020) (see Methods) (Supplementary Table 1 and Supplementary Fig. 2). A substantial proportion of genes associated with ageing were differentially expressed in the older flies, indicating that the age at which we extracted the RNA was sufficient to classify them as old (see Methods) (Supplementary Fig. 3 and 4).

We used TEtools to assess whether TE expression varies with age, irrespective of sex (see Methods) (Supplementary Tables 3 and 4) (Lerat, et al. 2017). To do that, we first looked at the percentage of reads mapping to TE sequences in our RNA-seq data. Using a generalized linear model (GLM) analysis, we found that the percentage of TE-derived reads increase during ageing in all the species tested (estimate 0.314, p < 0.001) (Fig. 2A). In particular, *D. melanogaster* showed a greater difference between young and old flies compared to the other species (Fig. 2A). Furthermore, comparing to *D. melanogaster*, *D. simulans* exhibited less TE-derived reads (estimate -0.652, p < 0.001), whereas *D. suzukii* showed more (estimate 0.725, p < 0.001), consistent with their respective TE contents (Fig. 2A).

**Figure 2.**
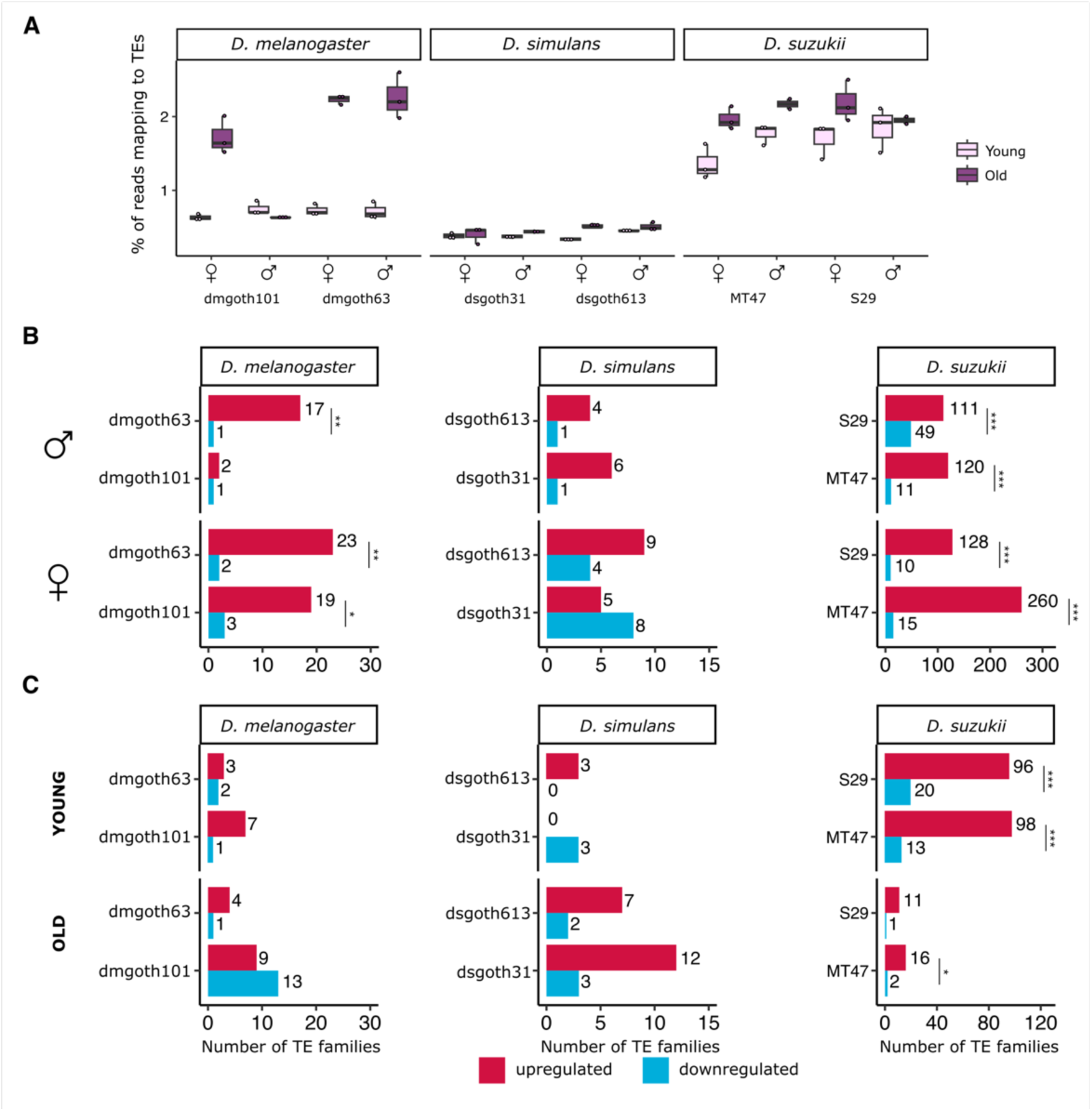
Ageing increases the number of TE transcripts and upregulated TE families in different *Drosophila* species. **A)** Boxplots showing the percentage of reads mapping to TE sequences in the RNA-seq data in young and old male and female flies of each strain. Boxplots show the median (horizontal line), first and third quartiles (lower and upper bounds, respectively), and minimum and maximum values (lower and upper whiskers, respectively). Each dot represents an independent biological replicate. **B**) Barplots showing the number of upregulated (in red) and downregulated (in blue) TE families with ageing for both males (upper panel) and females (bottom panel) in each strain. **C**) Barplots showing the number of upregulated (in red) and downregulated (in blue) TE families in young males compared to young females (upper panel) and in old males compared to old females (bottom panel) in each strain. Asterisks indicate level of statistical significance of chi-square tests (* p ≤ 0.05, ** p ≤ 0.01, ***p ≤ 0.001).

Since certain TE families might contribute more to the age-related increase in expression, we also analyzed TE expression at the family level. In both *D. melanogaster* and *D. suzukii*, we observed a greater number of upregulated TE families compared to downregulated ones in older flies across all strains (chi-square, all p < 0.05), except for males from the *D. melanogaster* dmgoth101 strain (chi-square, p = 0.999) (Fig. 2B). Similar results were found for both males and females, suggesting that TEs can be activated in ageing as it was observed in other studies from *D. melanogaster* and other species, such as humans (Fig. 2B) (Driver and McKechnie 1992; Nikitin and Woodruff 1995; Woodruff and Nikitin 1995; Li, et al. 2013; Van Meter, et al. 2014; Chen, et al. 2016; Wood, et al. 2016; Bogu, et al. 2019; Brown, et al. 2020b; Fabian, et al. 2021; Nguyen and Bachtrog 2021; Rigal, et al. 2022; Yang, et al. 2022; Teoli, et al. 2024; Tsai, et al. 2024). However, in *D. simulans*, the number of upregulated TE families did not statistically differ from the number of downregulated ones in older flies (chi-square, all p > 0.05), probably due to its reduced TE content. Nonetheless, there was a tendency towards a higher number of upregulated TE families (Fig. 2B). Although the number of upregulated TE families generally increases with age, these TE families showed little to no overlap between sexes or between strains of the same species (Supplementary Fig. 5). This result suggested that TE expression during ageing is not family specific.

Overall, we observed an increased number of upregulated TE families in older flies of *D. melanogaster* and *D. suzukii*. In *D. simulans*, there was a similar tendency, although it was not statistically significant, likely attributed to its reduced TE content. These findings are in line with the fact that ageing might be associated with the disruption of the epigenetic mechanisms that silence TEs favouring their activation.

### The age-related increase in TE-derived reads and expressed TE families is not influenced by sex

As flies age, the high TE content on the Y chromosome may contribute to higher TE expression in older males compared to females. We investigated this by comparing the percentage of TE-derived reads in older males and females.

Performing a GLM analysis, we found no statistically significant effects of sex (estimate 0.056, p = 0.398) or the interaction between sex and age (estimate -0.140, p = 0.271) on the percentage of TE-derived reads (Fig. 2A). Similarly, at the TE family level, we found no statistically significant differences in the number of upregulated and downregulated TE families between older males and older females, except in the *D. suzukii* MT47 strain (chi-square, p = 0.029) (Fig. 2C). However, there was a general trend toward finding more upregulated TE families (Fig. 2C). Again, little overlap between differentially expressed TE families were found between strains from the same species, suggesting that there are no particular TE families leading to the sex differences in TE expression of older flies (Supplementary Fig. 5). When comparing the expression of TE families between young male and female flies, we also observed no statistically significant differences between the number of upregulated and downregulated TE families in *D. melanogaster* and *D. simulans* (chi-square, p values > 0.05) (Fig. 2C). Conversely, in *D. suzukii*, we did observe more upregulated TE families in young males compared to young female flies (chi-square, p values < 0.001) (Fig. 2C).

Altogether, we found no significant sex differences in the overall age-related increase of TE transcripts, nor in the number of TE families up-or downregulated in older males versus older females.

### *D. melanogaster* and *D. suzukii* flies show higher abundance of expressed TE copies with age

We previously observed that TE-derived reads, as well as some TE families, increase with age, without apparent sex-specific effects. However, our family-level approach might mask differences in the expression of individual TE copies, which could be selectively activated or released due to locus-specific chromatin relaxation. To address this, we used Telescope to precisely measure TE expression at specific genomic locations and assess the impact of age and sex on individual TE copies (see Methods) (Bendall, et al. 2019) (Supplementary Tables 3 and 4). Telescope is a computational tool that resolves ambiguous mapped reads by probabilistically assigning them to their most likely source transcript using a Bayesian model (Bendall, et al. 2019). We performed this analysis with three biological replicates to ensure the robustness and reliability of our results.

On the one hand, we found a general trend towards more upregulated than downregulated TE copies with age in both males and females of *D. melanogaster* and *D. suzukii* (Fig. 3A). These results were statistically significant (chi-square, all p < 0.05), except for the *D. melanogaster* females of the dmgoth101 strain and the *D. suzukii* males of the S29 strain (chi-square, all p > 0.05) (Fig. 3A). On the other hand, in *D. simulans*, the number of upregulated TE copies did not statistically differ from the number of downregulated ones (Fig. 3A). These findings are in line with our previous analysis on TE-derived reads analysed with TEtools software (Fig. 2A). Moreover, as expected based on the TE content, *D. suzukii* showed an increased number of differentially expressed TE copies (DETEs) in both males and females compared to the other species (Fig. 3A).

**Figure 3.**
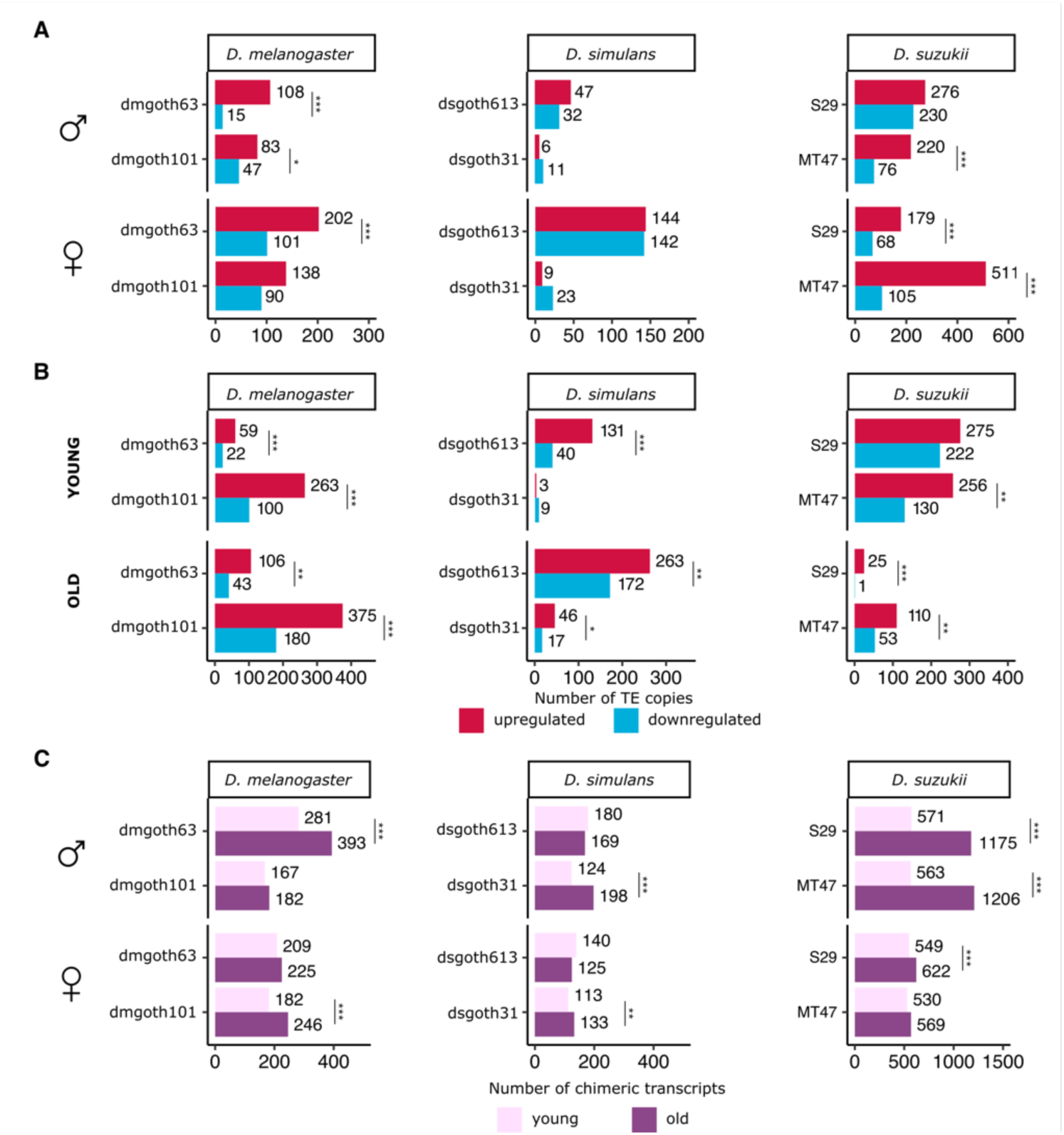
Ageing increases the number of activated TE copies in different *Drosophila* species. **A**) Barplots showing the number of upregulated (in red) and downregulated (in blue) TE copies with ageing for both males (upper panel) and females (bottom panel) in each strain. **B**) Barplots showing the number of upregulated (in red) and downregulated (in blue) TE copies in young males compared to young females (upper panel) and in old males compared to old females (bottom panel) in each strain. **C**) Ageing increases the generation of chimeric transcripts in some samples. Barplots showing the number of identified chimeric transcripts per strain as a sum of chimeric transcripts supported only by paired-end reads that have mapped in both transcripts and TE sequences (“chimreads_evidence”), chimeric transcripts supported only by the transcriptome assembly (“transcriptome_evidence”), and chimeric transcripts for which both previous methods have predicted the same chimeric transcripts (“double_evidence”). Asterisks indicate level of statistical significance of chi-square tests (* p ≤ 0.05, ** p ≤ 0.01, ***p ≤ 0.001).

We also tested whether male and female flies from the same strain shared some of the DETEs. Indeed, we found that some of the upregulated DETEs in males were shared with females (Supplementary Fig. 6). These shared DETEs are therefore absent from the Y chromosome, indicating that a subset of activated TEs might not be present in the Y chromosome, at least in some strains, suggesting a broader, global influence of the Y chromosome on TE expression. Overall, we observed an increased number of upregulated TE copies in older flies of *D. melanogaster* and *D. suzukii*.

### More specific TE copies are expressed in older males compared to older females

Using a copy-specific TE expression approach, we tested whether the higher TE abundance in males, driven by their TE-rich Y chromosome, results in a greater number of expressed TE copies in aging males compared to aging females.

We detected more upregulated TE copies than downregulated ones in old male flies compared to old female flies in all the strains of the three species tested (chi-square, all p < 0.05) (Fig. 3B). These differences are already observed at young ages (chi-square, all p < 0.05), although it is not the case for the *D. simulans* dsgoth31 and the *D. suzukii* S29 strains (chi-square, p = 0.399 and p = 0.105, respectively) (Fig. 3B). In contrast to the other two species, *D. suzukii* showed a greater number of DETEs between sexes in young flies than in old flies (Fig. 3B).

We also found that the upregulated TE copies observed between old males and females are present in all chromosome arms, with an expected concentration in chromosome centromeres and in the dot chromosome (Supplementary Fig. 7). In all strains, full-length TE copies, which are more likely to transpose, were significantly overrepresented among upregulated TEs compared to their overall genome proportion (chi-square, p < 0.001). This suggested that age-dependent TE expression may lead to new insertions; however further experiments are needed to confirm this.

We finally performed a permutation test to check whether DETEs found in old male flies are enriched nearby (+-1kb) differentially expressed genes (DEGs) (see Methods). Overall, we found no differences (p-values > 0.05) or statistically significant depletion (p-values < 0.05) in both upregulated and downregulated TE insertions nearby genes (Supplementary Table 5).

In summary, using a copy-specific approach, comparisons between older male and older female flies revealed an increased number of expressed TE copies than repressed. The copy-specific TE expression analysis revealed sex differences that were likely masked in family-level analysis.

### Ageing increases the number of TE chimeric transcripts in *D. suzukii* males

TEs can influence nearby genes by altering their expression and structure, potentially generating chimeric transcripts with both gene and TE sequences (Casacuberta and González 2013; Coronado-Zamora and González 2023; Oliveira, et al. 2023). To investigate if age-dependent TE activation is linked to increased chimeric transcripts, we used the ChimeraTE pipeline to quantify gene-TE chimeric transcripts in RNA-seq data from young and old flies of *D. melanogaster*, *D. simulans*, and *D. suzukii* (Oliveira, et al. 2023). We define the number of gene-TE chimeric transcripts as the count of distinct chimeric transcripts (isoforms) found in the dataset, rather than the number of reads mapping to these chimeric transcripts.

Older flies showed a significant increase in chimeric transcripts compared to younger flies in some samples: dmgoth63 males, dmgoth101 females, dsgoth31 males and females, S29 males and females, and MT47 males (chi-square, all p < 0.001) (Fig. 3C). Remarkably, in *D. suzukii*, males exhibited over twice as many chimeric transcripts in older flies compared to younger ones (Fig. 3C). Moreover, within each strain, there is only a small proportion of shared chimeric transcripts between different sexes or age groups (Supplementary Fig. 8).

In all species, there was a moderate positive correlation between chimeric transcript generation and TE family copy number (Pearson correlation all p < 0.05) (Supplementary Fig. 9). In *D. melanogaster*, the TE families *roo*, *INE-1*, and *copia* were most frequently associated with chimeric transcripts, with *roo* and INE-1 being abundant in both young and old flies, and *copia* showing increased chimeric transcript formation in older flies (Coronado-Zamora and González 2023; Oliveira, et al. 2023) (Supplementary Fig. 10). In *D. simulans*, the most prevalent families producing chimeric transcripts were *roo*, *INE-1*, *P-elements*, and *GATE* elements, whereas in *D. suzukii*, it was predominantly *Jockey-3* and *TART* elements (Supplementary Fig. 11 and 12).

Finally, we quantified the proportion of genes producing chimeric transcripts that were found differentially expressed with age. The contribution of these genes was very low: in *D. melanogaster* we found on average 2.92% in males and 5.34% in females; in *D. simulans*, 1.64% in males and 6.21% in females; and in *D. suzukii*, 4.75% in males and 3.56% in females. Moreover, there was no enrichment of DEGs among those generating chimeric transcripts in any sample (chi-square tests, all p > 0.05). Gene-TE chimeric transcripts could interfere with TE quantification. If a highly expressed host gene produces a chimeric transcript containing a TE sequence, part of the reads from that transcript could be misassigned to the TE, thereby artificially inflating its estimated expression level. However, although gene–TE chimeric transcripts could in theory inflate TE expression, their contribution in our data is very limited, and we do not expect them to affect our overall results.

Altogether, we observed an increase in gene-TE chimeric transcripts with age in some samples, particularly in *D. suzukii* males. While these changes may relate to age-dependent TE activation, they could also result from increased gene expression or reduced splicing efficiency with age (Wood, et al. 2016; Bhadra, et al. 2020).

### Increased TE expression in aged males could be associated with reduced lifespan both *D. melanogaster* and *D. simulans*

To investigate whether the increased expression of some specific TE copies in aged males compared to aged females correlates with sex lifespan differences, we performed longevity assays across all strains from our transcriptome analysis.

Males showed reduced lifespan in *D. melanogaster* and *D. simulans* (Log Rank all p < 0.001), with negative SGL values (Fig. 4A-D, and Supplementary Table 6). Nevertheless, in *D. suzukii*, differences in lifespan between males and females were not statistically significant (Log Rank p = 0.400 and p = 0.500 for MT47 and S29, respectively), though negative SGL values suggest some variance in ageing associated with sex (Fig. 4E and 4F and Supplementary Table 6). Thus, the increased number of DETEs in aged males compared to aged females (Fig. 3B) might be associated with reduced lifespan in *D. melanogaster* and *D. simulans*, but not clearly in *D. suzukii* (Fig. 4).

**Figure 4.**
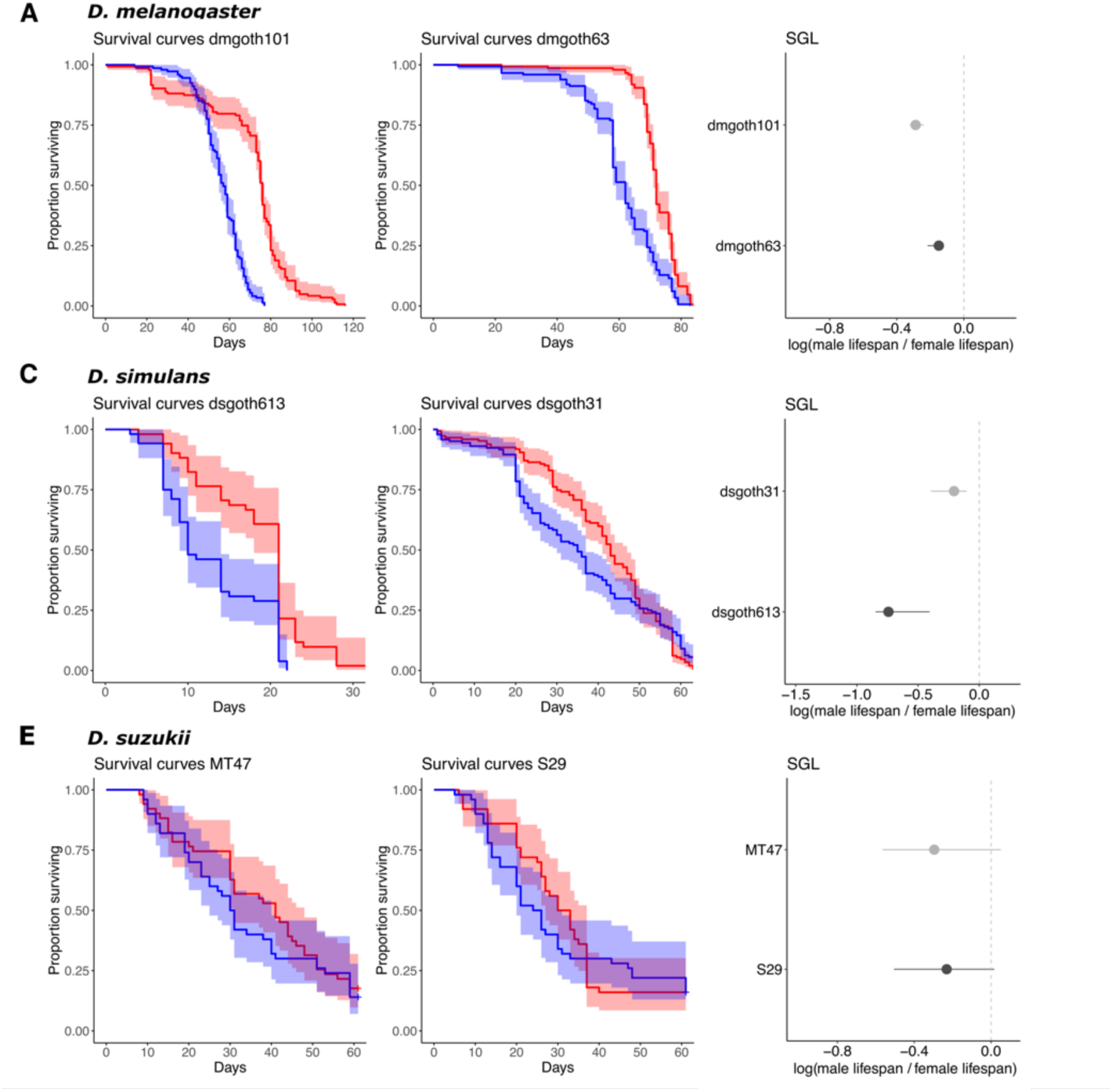
*D. melanogaster* and *D. simulans* males have reduced lifespan. Kaplan–Meier survivorship curves for males (blue) and females (red) in **A**) *D. melanogaster*, **C**) *D. simulans,* and **E**) *D. suzukii* with the shaded region indicating the upper and lower 95% confidence interval calculated from the Kaplan–Meier curves. SGL values measured as the logarithm of the ratio between male and female median lifespan are shown for each strain of **B**) *D. melanogaster*, **D**) *D. simulans,* and **F**) *D. suzukii*. Dots represent mean SGL values and lower and upper whiskers indicate the upper and lower 95% confidence intervals.

Finally, when analyzing each strain individually, we found no correlation between male lifespan or SGL values and their overall TE content (Pearson correlation all p < 0.05). Hence, overall TE content does not appear to be directly linked to reduced lifespan, as previously suggested for *D. melanogaster* and other species like the Greenland shark (Delanoue, et al. 2023; Sahm, et al. 2024).

### Y chromosome replacement leads to increased SGL by reducing male lifespan

To check whether the Y chromosome is associated with changes in SGL, we performed introgression experiments to replace the Y chromosome between *D. melanogaster* strains with different SGL values (see Methods).

Introgression crosses were made between strains dmgoth101 (−0.29 SGL, 95% CI [-0.32, - 0.24]) and dmsj7 (−0.07 SGL, 95% CI [-0.13, -0.04]), with longevity assays conducted over ten generations. “A crosses” started with dmgoth101 females and dmsj7 males, while “B crosses” were the reciprocal. Both crosses showed an increase in SGL values across generations, primarily due to a reduction in male lifespan (Fig. 5A and Supplementary Table 7). In A crosses, SGL rose from -0.26 (95% CI [-0.35, -0.19]) to -1.11 (95% CI [-1.20, -1.01]) from generation F1 to F10 (Fig. 5A). Similarly, B crosses showed an initial rise from -0.23 (95% CI [-0.29, -0.14] to -0.68 (95% CI [-0.71, -0.60]) by F5 but decreased by F10 (95% CI [-0.27, - 0.01])) (Fig. 5A). In both reciprocal crosses, males exhibited a progressive reduction in lifespan across all generations, unlike females, whose lifespan showed less variation across generations (standard deviation of mean lifespan in A crosses: 22.21 for males and 12.13 for females; B crosses: 14.42 for males and 10.06 for females) (Fig. 5B and 5C). Repeating the experiment with strains that did not show SGL differences (dmgoth101 and dmgoth63) did not result in significant SGL value increases across generations (Supplementary Fig. 13 and Supplementary Table 7).

**Figure 5.**
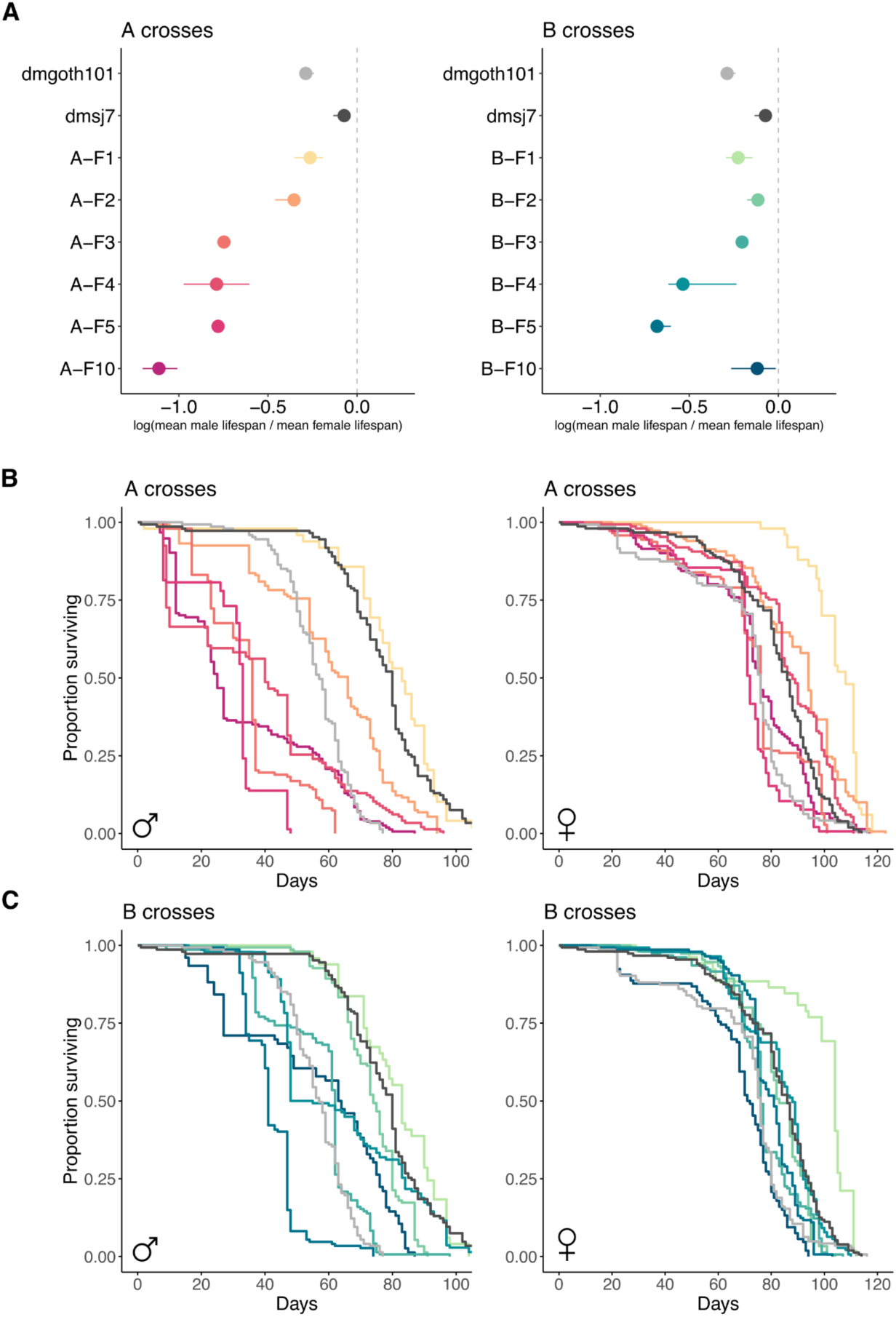
Y chromosome replacement is associated with reduced male lifespan. **A**) SGL values measured as the logarithm of the ratio between male and female median lifespan are shown for A reciprocal crosses (left panel) and B reciprocal crosses (right panel) of *D. melanogaster*. Results for maternal strains are shown in grey (dmgoth101) and black (dmsj7) while different colours represent generations of introgression from F1 to F10. Dots show mean SGL values and lower and upper whiskers indicate the upper and lower 95% confidence intervals. **B**) Kaplan–Meier survivorship curves for males (left panel, n=150) and females (right panel, n=150) in A reciprocal crosses. **C**) Kaplan–Meier survivorship curves for males (left panel, n=150) and females (right panel, n=150) in B reciprocal crosses. Male lifespan between dmgoth101 and dmgoth101 was statistically significantly different (Log-Rank test p < 0.001).

In summary, we found that the replacement of the Y chromosome between strains with different SGL values is associated with a progressive increase in SGL due to a reduced male lifespan. If SGL values were directly linked to the Y chromosome’s TE content, we might expect the replacement of the Y chromosome to result in an SGL value matching that of the donor strain across generations. However, this was not observed since SGL values increased in both reciprocal crosses when replacing the Y chromosome between strains with different SGL values. While further experiments are needed, these findings indicate that the modification of the TE composition and distribution on the Y chromosome and/or the Y chromosome as a whole may have a negative impact in male lifespan.

### Y chromosome replacement is associated with an increase TE expression

To assess the impact of Y chromosome replacement on TE expression, we extracted RNA from 5-7 days old male and female *D. melanogaster* flies at generations F1 and F10 in reciprocal introgression crosses between dmgoth101 and dmsj7 (see Methods). TE expression was measured using TEtools (Lerat, et al. 2017).

We assessed the effects of reciprocal cross (A or B), generation (F1 or F10), sex, and their interactions on TE-derived reads percentage (see Methods). F10 flies had more TE-derived reads than F1 in both sexes (GLM, estimate 0.105, p = 0.033), with differences more pronounced in B crosses (GLM, estimate 0.068, p = 0.048) (Fig. 6). In addition, males had more TE-derived reads than females in both generations (GLM, estimate 0.295, p < 0.001), and the interaction between generation and sex was not statistically significant (GLM, estimate 0.077, p = 0.249) (Fig. 6). These findings suggested that the Y replacement might be associated with increased TE transcripts across generations.

**Figure 6.**
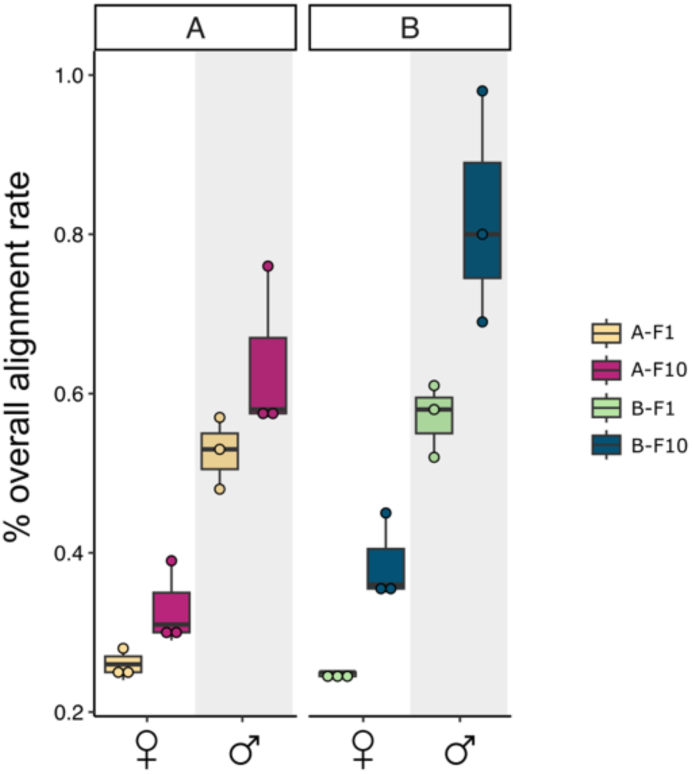
Y chromosome replacement is associated with increased TE expression. Boxplots showing the percentage of reads mapping to TE sequences in the RNA-seq data in F1 and F10 generations of introgression in both A and B reciprocal crosses. Boxplots show the median (horizontal line), first and third quartiles (lower and upper bounds, respectively), and minimum and maximum values (lower and upper whiskers, respectively). Each dot represents an independent biological replicate.

Overall, Y chromosome replacement between strains with different SGL values was associated with increased TE expression in males compared to females already in the first introgression cross (F1). The percentage of TE-derived reads increased in F10 compared to F1 in both sexes and reciprocal crosses.

## DISCUSSION

In this work, we characterized SGL variability across wild-type populations in three *Drosophila* species, finding greater population-level variance and an overall reduced male lifespan compared to females, consistent with previous studies (Yoon and Gagen 1990).

Sex-differences in lifespan have been associated with TE expression in some organisms such as *D. melanogaster* and *D. miranda* (Brown, et al. 2020b; Nguyen, et al. 2022). Here, using a copy-specific approach, we found a general age-related increase in TE copies in both males and females of *D. melanogaster* and *D. suzukii*. This result suggests that there might be an age-related derepression of TEs, likely due to a general loss of heterochromatin in repetitive regions in aged flies, as it was previously observed in *D. melanogaster* (Brown, et al. 2020b). Specifically, male flies from the same species have been shown to undergo a more rapid heterochromatin mark loss producing a more pronounced derepression of TEs in older males compared to older females (Brown, et al. 2020b).

Previous studies in a *D. melanogaster* laboratory strain showed some differences in the total fraction of repetitive reads that increased during ageing between males (4.2%) and females (2.6%) (Brown, et al. 2020b). Here, we could not detect statistically significant effects of sex or the interaction between sex and age on the percentage of TE-derived reads, indicating that age is associated with a similar increase in TE transcript levels in both males and females. In this work, the difference in fraction of TE-derived reads between old and young samples was 1.47% in males and 1.37% in females from dmgoth63; and 0.07% in males and 0.01% in females from dsgoth31. Moreover, it was previously found in a laboratory strain that 32 TE families showed a significant increase in expression during ageing in males while only six were found in females (Brown, et al. 2020b). Although we did not follow the same approach, differential expression analysis between older males and older females did reveal more upregulated specific TE copies in older males in all the species tested. The TE expression analysis at the copy-level allowed to uncover sex differences that were likely diluted in family-level assessments. These results suggested that, although TE expression should be seen as an indirect evidence of genomic instability, male genome integrity might be more susceptible to ageing effects compared to females.

Indeed, particularly in *D. melanogaster* and *D. simulans* species, we found that increased TE copy expression in aged males compared to aged females might be associated with reduced lifespan. However, we could not find a significant correlation between male lifespan or SGL values and overall TE content. This paradox has been explored in long-lived species, which often exhibit an overall reduction in both TE abundance and expression (Elsner, et al. 2018; Ricci, et al. 2023; Sturm, et al. 2023; Teefy, et al. 2023). However, exceptions exist where TEs might play a beneficial role in ageing (Sulak, et al. 2016; Zhao, et al. 2021; Sahm, et al. 2024). For instance, in blind mole rats (*Spalax galili*), TE activation can act as a tumour suppressor by triggering the innate immune system (Zhao, et al. 2021). In the Greenland shark (*Somniosus microcephalus*), the longest-lived vertebrate, TEs constitute the 70.6% of the genome, the highest proportion described in sharks (Sahm, et al. 2024). In this case, TE expansion may be linked to the expansion of DNA repair genes, which in turn may help to tolerate higher levels of TE activity (Sahm, et al. 2024). Therefore, unraveling the complex mechanisms of delayed aging requires careful consideration of species-specific features, including the abundance, composition, and distribution of TE in the genomes (Yushkova and Moskalev 2023).

Besides TE expression, it has been proposed that age might increase TE transposition, leading to a greater number of somatic TE insertions that impact genomic stability and cellular homeostasis, ultimately contributing to a decreased lifespan. In *D. melanogaster*, it has been shown that lifespan is decreased as somatic genetic damage produced by *P element*, *mariner*, and *mdg4* insertions (Nikitin and Woodruff 1995; Woodruff and Nikitin 1995; Li, et al. 2013; Wood, et al. 2016; Chang and Dubnau 2019). Yet, some recent studies found that the number of new TE insertions do not seem to correlate with age, suggesting that TE expression instead could have a global impact in transcriptional dysfunction during ageing and affecting lifespan (Treiber and Waddell 2017; Rigal, et al. 2022; Yang, et al. 2022; Pabis, et al. 2024; Tsai, et al. 2024). Here, we observed that ageing increases the generation of TE-chimeric transcripts in some samples, and specially in *D. suzukii* males. Thus, our findings suggest that age-related TE expression could influence transcript structure by increasing the presence of TE-chimeric transcripts in *D. suzukii* males, hinting that the impact on fly lifespan may not solely be attributed to TE somatic transposition.

Our results also revealed that, in *D. melanogaster,* we can manipulate SGL values. The replacement of the Y chromosome between strains exhibiting differences in SGL values and male lifespan was associated with a progressively decline in male survival, leading to a more pronounced SGL. Further analysis will be needed to determine whether Y chromosome size differences (associated with a different Y-linked TE abundance) or differences in TE distribution are responsible for the observed effects. In fact, two recent works indicate that male survival is not linked to Y chromosome size in both *D. melanogaster* and *D. pseudoobscura* (Nguyen, et al. 2022; Delanoue, et al. 2023). Hence, based on our introgression experiments, we hypothesize that differences in TE composition among Y chromosomes (proportion of full-length copies or average enrichment in repressive histone marks) may offer a more plausible explanation for variances in male lifespan compared to Y chromosome sizes or TE abundance (Lemos, et al. 2010; Griffin, et al. 2015). However, it is also conceivable that other genomic regions other than TEs on the Y chromosome are contributing to reduced lifespan, as well as mitochondrial genome – Y chromosome interactions (Ågren, et al. 2020). Thus, while TEs are strong candidates, we cannot conclude beyond the fact that some factor on the Y chromosome is linked to reduced lifespan.

In summary, this work reveals that SGL is a variable trait in the wild, and emphasizes the importance of studying the role of TEs in the evolution of sex differences in lifespan across different species.

## METHODS

### Fly strains and rearing

Samples of wild-type populations of *D. melanogaster*, *D. simulans,* and *D. suzukii* were collected, and different isofemale strains were established for each species and geographical origin directly from gravid females from the field. A comprehensive list of all populations along with collection information is available in Supplementary Table 1. *D. melanogaster* and *D. simulans* fly strains were reared in vials with a nutritive “LM” medium containing agar, maize flour, yeast, Nipagine, ethanol and water, while *D. suzukii* strains were reared in vials with “Dalton” medium containing agar, cornmeal, yeast, sugar, Nipagine, ethanol and water. Fly strains were kept in a 12:12 hour light/dark cycle at 25 °C.

### Longevity assays

#### Longevity assays for the analysis of SGL natural variation in different Drosophila species

50 recently (less than 8 hours) emerged males and 50 virgin females from each isofemale strain were placed in groups of 10 in tubes containing nutritive medium (“LM” for *D. melanogaster* and *D. simulans* and “Dalton” for *D. suzukii*). Each tube was considered as an independent biological replicate thus having five replicates per sex and strain. Flies were separated by sex without anaesthesia using an aspirator. Dead flies in every vial were counted every day, except for weekends, until 80% of the flies in the tube were dead. Alive flies were placed in new tubes with fresh medium every week. Longevity assays were performed at 25°C and at constant humidity of 60%.

#### Longevity assays in the inbred strains included in the transcriptome analysis

For *D. melanogaster* strains, 150 recently (less than 8 hours) emerged males and 150 virgin females from each strain were placed in groups of 10 in tubes containing “LM” medium. Each tube was considered as an independent biological replicate thus having 15 replicates per sex and strain. For *D. simulans* and *D. suzukii* strains, 50 recently (less than 8 hours) emerged males and 50 virgin females from each strain were placed in groups of 10 in tubes containing nutritive medium (“LM” for *D. simulans* and “Dalton” for *D. suzukii*). Each tube was considered as an independent biological replicate thus having five replicates per sex and strain. Dead flies in every vial were counted every day, except for weekends, until 80% of the flies in the tube were dead. Alive flies were placed in new tubes with fresh medium every week. Longevity assays were performed at 25°C and at constant humidity of 60%.

#### Longevity assays in the Y introgressed strains

150 recently (less than 8 hours) emerged males and 150 virgin females from each strain/generation of introgression were placed in groups of 10 in tubes containing “LM” medium. Again, each tube was considered as an independent biological replicate thus having 15 replicates per sex and strain/generation of introgression. Flies were separated by sex without anaesthesia using an aspirator. Dead flies were counted every day, except for weekends, until all the flies in the tubes were dead. Alive flies were placed in new tubes with fresh medium every week. Longevity assays were performed at 25°C and at constant humidity of 60%.

### SGL values calculation

We estimated sex-specific median adult lifespan (in days) for each strain/generation of introgression in our dataset (Supplementary Tables 2, 6, and 7). The median adult lifespan corresponds to the time when 50% of the individuals in a particular strain are dead. To estimate the confidence interval for both males and females of each strain, we implemented a bootstrapping procedure. We performed 100,000 bootstrap iterations to calculate the median adult lifespan for each sample and then we calculated the logarithm ratio (ln, base e) of median longevity between males and females (SGL values) (Lemaître, Ronget, Tidière, et al. 2020). Confidence intervals for the difference in median longevity between males and females was computed using quantiles of the bootstrap of the logarithm ratios.

For the analysis of SGL natural variation, we used a Bayesian approach implemented in the MCMCglmm package in R to partition the variance and identify the contribution of species, population, and sampling variation to the observed differences in SGL. We fitted a hierarchical regression model including species as a fixed effect and population as a random effect, with SE weighting for the longevity measurements. The priors were set as follows: for the residual variance, we used an inverse-Wishart prior with nu = 0.002; for the population random effect, we used an inverse-Wishart prior with nu = 1 and a large variance (a = 1000) for the prior mean (diff_median_longevity∼species+(1∣population)+idh(SE):units).

We also performed survival analysis using the Kaplan-Meier method and Cox proportional hazard models. The survival analysis was conducted separately for male and female flies within each strain/generation of introgression. We used the *survfit* function in R to estimate the survival probabilities over time based on the event status (dead or alive). Additionally, we generated survival curves using the *ggsurvplot* function from the *survminer* R package. These curves represent the estimated survival probabilities over time using a Kaplan-Meier estimation. Finally, we conducted pairwise comparisons to evaluate the significance of difference in survival distributions among various samples. This analysis was performed using the *pairwise_survdiff* function, applying the Benjamini-Hochberg correction method for multiple comparisons.

### DNA extraction, nanopore sequencing and genome assemblies

Genome assemblies for *D. melanogaster* dmgoth101 and dmgoth63 and for *D. simulans* strains dsgoth31 and dsgoth613 were obtained from Mohamed et al., 2020 (Mohamed, et al. 2020). To obtain genome assemblies of *D. suzukii*, we sequenced with Oxford Nanopore Technologies (ONT) two inbred strains: S29 and MT47. DNA was extracted using phenol chloroform from a pool of 10 adult females. The quantity and quality of genomic DNA were assessed using a NanoDrop™ One UV-Vis spectrophotometer (Thermo Fisher Scientific) and a Qubit® 1.0 Fluorometer (Invitrogen). Three micrograms of DNA were then repaired with the NEBNext FFPE DNA Repair Mix (NEB M6630). End repair and dA-tailing were carried out using the NEBNext End Repair/dA-Tailing Module (E7546, NEB). Ligation was evaluated with the Ligation Sequencing Kit 1D. MinION sequencing was conducted according to the manufacturer’s instructions using R9.4.1 flow cells (FLO-MIN106, ONT) and a Nanopore MinION Mk1b sequencer (ONT), operated by ONT MinKNOW (v.18.3.1). Following sequencing, base calling was performed using *Guppy* (v.4.0.5) in high-accuracy mode.

Quality control of Nanopore reads was conducted using *NanoPlot* (v.1.10.2) (https://github.com/wdecoster/NanoPlot). Reads with a quality score below 7 were excluded from further analysis. Contig assembly was performed with *Flye* (v.2.8) (Heng Li 2013) using default parameters, except for the –plasmids option. All contigs were aligned with *minimap2* (v2.16) (Li 2018) using the –x map-ont option, followed by four rounds of polishing with *RACON* (v1.3.2) (Vaser, et al. 2017) under default settings. Assembly quality metrics were evaluated using *Assembly-Stats* (v1.0.1) (https://github.com/sanger-pathogens/assembly-stats). Any assembly inconsistencies were manually reviewed with *D-genies* (v1.2.0) (Cabanettes and Klopp 2018) and corrected using *samtools* (v1.9.0) with the *faidx* function, and *Gepard* (v1.4.0) was used to identify breaking points. Super scaffolding of all corrected assemblies was carried out with *RaGOO* (v.1.1), using the –s and –t 4 parameters, based on the reference genome assembly (Paris, et al. 2020). The resulting chromosome-scale assemblies were then analyzed using Benchmarking Universal Single-Copy Orthologs (BUSCO) (v.5.4.3) (Simão, et al. 2015) with the diptera_odb10 lineage (−l parameter). Gene annotation for the Nanopore genomes was completed with *Liftoff* (v.1.6.3), referencing the WT3_2.0 *D. suzukii* genome (Paris, et al. 2020).

### RNA-seq extraction and sequencing

RNA was extracted from six inbred strains, comprising two strains for species: *D. melanogaster* (dmgoth101 and dmgoth63), *D. simulans* (dsgoth31 and dsgoth613) and *D. suzukii* (MT47 and S29). For young samples, male and female flies of each strain were separately placed in several tubes with nutritive medium in groups of 20-25. When flies were 5 day-old, 30 flies per sex were dissected and ovaries and testes were removed from female and male flies, respectively, and only carcasses were kept. For old samples, male and female flies of each strain were also separately placed in several tubes with nutritive medium in groups of 20-25. 30 flies per sex were dissected and ovaries and testes were removed from female and male flies, respectively, and only carcasses were kept when at least 25% of the flies in the tube were dead. Consequently, older samples may exhibit varying chronological ages across strains. Three replicates per strain, age and sex were performed. Then, for both young and old samples RNA was extracted from the obtained carcasses. For the Y chromosome replacement experiments, RNA was extracted from both F1 and F10 male and female flies from the two reciprocal crosses. Samples were obtained as mentioned before when flies were 5 days-old.

RNA extraction was carried out using the RNeasy Mini kit (Qiagen) following manufacturer’s instructions and then a DNase treatment (Invitrogen) was performed. mRNA libraries were prepared with the Illumina TruSeq RNA Sample Prep Kit, following the manufacturer’s guidelines. Libraries were sequenced on the Illumina HiSeq 4000 platform with paired-end 100 nt reads. The sequence raw files have been deposited in NCBI SRA under the BioProject accession number PRJNA1159534.

### RNA-seq mapping and gene quantification

Quality of the *fastq* sequencing files was assessed with FastQC (v.0.11.9). Then, *fastp* (v.0.23.4) was used to filter out bad reads, adapter removal and trimming (Chen 2023). Trimmed reads from *D. melanogaster*, *D. simulans,* and *D. suzukii* were then mapped against the Flybase *D. melanogaster* (r6.16) (Öztürk-Çolak, et al. 2024), the *D. simulans* (r2.02) (Öztürk-Çolak, et al. 2024), and *D. suzukii* WT3_2.0 (Paris, et al. 2020) reference genomes, respectively with *STAR* (v.2.7.11a) (Dobin, et al. 2013) using default parameters. On average, 92.7% of the reads were uniquely mapped to the corresponding genomes. To quantify gene expression, we used *htseq* (v.2.0.3) (Anders, et al. 2015), with parameter *-t exon*, which uses the sum of the reads aligned in all exons to assign counts for each gene.

### TE annotation

All the six strains used in the transcriptomic analysis (dmgoth101, dmgoth63, dsgoth31, dsgoth613, MT47, and S29) have been sequenced with long-read sequencing technologies and genome assemblies were available or generated (Mohamed, et al. 2020; Mérel, et al. 2021). TE insertions were identified in the genome assemblies using *RepeatMasker* (v.4.1.2), which masks the genome where there are fragments homologous to consensus TE sequences. Parameters used were: *-cutoff 250, -a, -s, -norna*. Consensus TE sequences used for *D. melanogaster* was obtained by using the parameter *-species drosophila.* However, consensus sequences for *D. simulans* were obtained from https://github.com/bergmanlab/drosophila-transposons/, while for *D. suzukii* from https://github.com/vmerel/DSu-TE/tree/main/TE_db. Finally, *bed* files containing the coordinates of each annotated TE were created for every genome.

### Differential expression analysis of genes

Differential expression analyses of genes were conducted using the R package *DESeq2* (v.1.40.2) (Love, et al. 2014). Raw read counts were merged into a unified matrix for each strain and subsequently normalized using default parameters (Supplementary Tables 3 and 4). Differential expression of genes was assessed between old and young samples in each strain separately. The log2 fold-change was adjusted using the *apeglm* (v.1.22.1) algorithm from the *lfcshrink* function to minimize noise and preserve significant differences (Zhu, et al. 2019). Genes were considered as significantly differentially expressed when the adjusted p-value was < 0.05 and the log2 fold-change > |1|.

### Age-related genes found differentially expressed in old flies

Genes previously associated with ageing in *D. melanogaster* were obtained from the Animal Ageing and Longevity Database (AnAge) (https://genomics.senescence.info/genes/search.php?search.php?show=1&sort=1&page=1&organism=Drosophila%20melanogaster). From this list, only genes with a described “increased” lifespan effect were selected. Then, gene symbols were converted to FlyBase IDs with the *Batch Download* online tool (https://flybase.org/batchdownload). We finally looked for overlap of the age-associated genes and the DEGs found in the young versus old samples comparisons. For *D. simulans*, we used the gene ortholog table available in FlyBase to look for correspondences in gene IDs. We could analyze a total of 149, 152, and 120 age-associated genes in *D. melanogaster, D. simulans*, and *D. suzukii*, respectively.

### Quantification and differential expression analysis for TE families

TE quantification at a family level was performed with *TEtools*, using the TEcount module (Lerat, et al. 2017). One of the inputs of the TEcount module is the *rosette* file, a tabular file containing the names of each TE copy in the genome that easily groups them in TE families (or other criteria if needed) (Lerat, et al. 2017). To do that, for each strain, we first build a *fasta* file with all the TE sequences annotated using *bedtools getfasta* and the previously generated *bed* file with TE annotations. Then, we generated the *rosette* file writing in the first column the name of the specific TE copy and in the second column the TE family in which it belongs. For the TE quantification of the introgressed F1 and F10 generations, a merged *rosette* file between the strains dmgoth101 and dmsj7 was used. Differential expression analysis of TE families was conducted using the R package *DESeq2* (v.1.40.2) as mentioned before, and merging in a single matrix gene and TE family counts (Love, et al. 2014) (Supplementary tables 3 and 4). TE copies were considered as significantly differentially expressed when the adjusted p-value was < 0.05 and the | log2 fold-change | > 1.

### RNA-seq mapping, quantification and differential expression analysis for specific TE copies

Quantification of TE copies was performed using *Telescope* (Bendall, et al. 2019). Briefly, for each strain, trimmed paired-end reads were aligned to the corresponding genome with *bowtie2* (v.2.5.1) using the parameter *-k 100*. Then, again for each strain, we generated a *gtf* file containing all the TE copies present in the strain following the format specified by *Telescope*. After that, the generated *sam* file was converted to *bam* with *samtools view* (v.1.17). Finally, for the locus-specific quantification of TEs, the function *telescope assign* was used taking the already generated *gtf* and *bam* files as inputs. Differential expression analysis of TE copies was conducted using the R package *DESeq2* (v.1.40.2) as mentioned before, and merging in a single matrix gene and copy-specific TE counts (Love, et al. 2014) (Supplementary tables 3 and 4). TE copies were considered as significantly differentially expressed when the adjusted p-value was < 0.05 and the | log2 fold-change | > 1.

### Identification of full-length insertions

Identification of full-length insertion among the upregulated TE copies found between old males and females (adjusted p-value was < 0.05 and | log2 fold-change | > 1) was performed with a generated *python* script that detects as full-length copies the ones that are at least 80% of the length of the consensus sequence and with a Kimura divergence value < 20 (values obtained from the previously generated *RepeatMasker* output files). For *D. melanogaster* and *D. simulans*, TE consensus sequences were obtained from https://github.com/bergmanlab/drosophila-transposons/, while *D. suzukii* consensus sequences were obtained from Mérel et al., 2021 (Mérel, et al. 2021).

### Permutation tests

To test whether DETEs found in old male flies were enriched nearby DEGs (+-1kb), we first created *bed* files with the +-1kb flanking regions for these genes in each specific genome. *bed* files with the coordinates of all TE copies were also generated for each strain. Then, we performed permutation tests by intersecting TE positions with gene regions (+-1kb) using the R package *regioneR* (Gel, et al. 2016), parameter *randomization = resampleRegions*, with 1000 random samples. Hence, for every up and downregulated gene set of size *n*, the software determines the count of those containing TEs in the +-1kb flanking regions. Subsequently, it conducts 1000 random samples of n genes each to assess if the observed number of genes with TEs deviated from the expected count in a genome-wide context (Supplementary Table 5).

### Identification of chimeric transcripts

Gene-TE chimeric transcripts were detected with *ChimeraTE* Mode 2 for each strain, sex, and age separately (Oliveira, et al. 2023). We used Mode 2 (chimTE_mode2.py), which identifies transcripts without a reference genome, using the prediction of chimeras from fixed and polymorphic TEs because we focused on capturing all the possible new age-related generated chimeric transcripts. Briefly, stranded RNA-seq reads were mapped against its respective annotation of transcripts and TE insertions. We used the transcript annotation from the release 6.54 *dmel-all-transcript-r6.54.fasta* for *D. melanogaster* (https://ftp.flybase.net/genomes/Drosophila_melanogaster/dmel_r6.54_FB2023_05/fasta/dme <u>l-all-transcript-r6.54.fasta.gz</u>); from the release 2.02 *dsim-all-transcript-r2.02.fasta* for *D. simulans* (https://ftp.flybase.net/genomes/Drosophila_simulans/dsim_r2.02_FB2017_04/fasta/dsim-all-transcript-r2.02.fasta.gz), and the *dsuz-all-transcripts-LBDM_Dsuz_2.1.fasta* file for *D. suzukii*(Paris, et al. 2020). Reference TE file for *D. melanogaster* and *D. simulans* was available online (https://github.com/bergmanlab/drosophila-transposons/tree/master/current), while reference TE file for *D. suzukii* was obtained online (https://github.com/vmerel/DSu-TE/tree/main/TE_db). We used the parameters *–strand rf-stranded* and *–assembly* to search for chimeric transcripts in the *de novo* transcript assembly. For each strain, sex and age, we got three output files: list of chimeric transcripts supported only by paired-end reads that have mapped in both transcripts and TE sequences, list of chimeric transcripts supported only by the transcriptome assembly, and chimeric transcripts supported by both mapped paired-end reads and transcriptome assembly. These three lists were merged and just the name of the gene and the TE family were kept to easily analyze the number of chimeric transcripts in each sample (Supplementary Table 8).

### Y chromosome replacement crosses

To check whether the Y chromosome is associated with changes in SGL, we performed introgression experiments to replace the Y chromosome between strains with different SGL values. The lack of recombination in *Drosophila* males enabled the generation of flies that differ only in the Y chromosome. Reciprocal crosses between the *D. melanogaster* strains dmgoth101 and dmsj7 were performed. We assigned the name “A crosses” to crosses initiated by mating a dmgoth101 female with a dmsj7 male, followed by successive backcrossing of male offspring with dmgoth101 females over ten generations. Conversely, “B crosses” represented the reciprocal pairing wherein dmsj7 females were initially mated with dmgoth101 males, followed by analogous backcrossing of male offspring with dmsj7 females. Five virgin females and five males aged 4-5 days old were placed in a tube containing a nutritive medium, with five replicates performed for each reciprocal cross. Every 2-3 days, the flies were transferred to new tubes with fresh medium until four tubes were obtained for each reciprocal cross. This ensured enough offspring to carry out longevity assays and facilitated the continuation of introgression crosses. F1 male flies were subsequently crossed with the maternal strain, and the male offspring of each generation was consistently crossed with the maternal strain for 10 generations. In each generation (except for generations F6-F9), longevity assays were carried out as previously described with 10 virgin females and 10 males from each of the five replicated reciprocal crosses (n=150).

### TE transcript quantification in introgression crosses

TE transcript quantification was performed with TEtools, using the TEcount module (Lerat, et al. 2017). One of the inputs of the TEcount module is the *rosette* file, a tabular file containing the names of each TE copy in the genome that easily groups them in TE families (or other criteria if needed) (Lerat, et al. 2017). To do that, for each strain (dmgoth101 and dmgoth63), we first build a *fasta* file with all the TE sequences annotated using *bedtools getfasta* and the previously generated *bed* file with TE annotations. Then, we generated a *rosette* file per strain writing in the first column the name of the specific TE copy and in the second column the TE family in which it belongs. After that, we merged the two *rosette* files. A GLM analysis was performed to assess the effects of the reciprocal cross, generation, sex, and the interaction between generation and sex on the percentage of TE-derived reads in R (glm(alignment_rate ∼ cross + generation + sex + generation:sex, family = gaussian(link = “identity”). The model showed a good fit (residual deviance = 0.1188, AIC = -47.315).

## DATA ACCESS

The sequencing data generated in this study has been submitted to the NCBI BioProject database (https://www.ncbi.nlm.nih.gov/bioproject/) under accession number PRJNA1159534.

Any additional information required to reanalyze the data reported in this paper is available from the lead contact upon request.

## ACKNOWLEDGEMENTS

This work was supported by the Agence Nationale de la Recherche (project LongevitY, grant Projet-ANR-20-CE02-0015), the GeEpiAdaptation project funded by the Marie Sklodowska-Curie Actions (project number 101065313), and the Fondation pour la Recherche Médicale. This work was performed using the computing facilities of the CC LBBE/PRABI. We also thank Marina Stamenkovic, Katja Hoedjes, Maaria Kankare and Darren Obbard for kindly providing some fly populations.

