## Supplementary Figures for "Exploring the role of transposable elements to sex gap in longevity in *Drosophila* species"

**Figure S1.** Natural variability in SGL in different *Drosophila* species. Histograms displaying the distribution of SGL values calculated as the logarithm of the ratio between male and female median lifespan. **A**) Histogram displaying the distribution of SGL values per population in *D. melanogaster*, *D. simulans*, and *D. suzukii*. UK: United Kingdom, FR: France, FI: Finland, NL: the Netherlands, BR: Brazil, ES: Spain, RS: Serbia. **B**) Histogram displaying the distribution of SGL values per species.

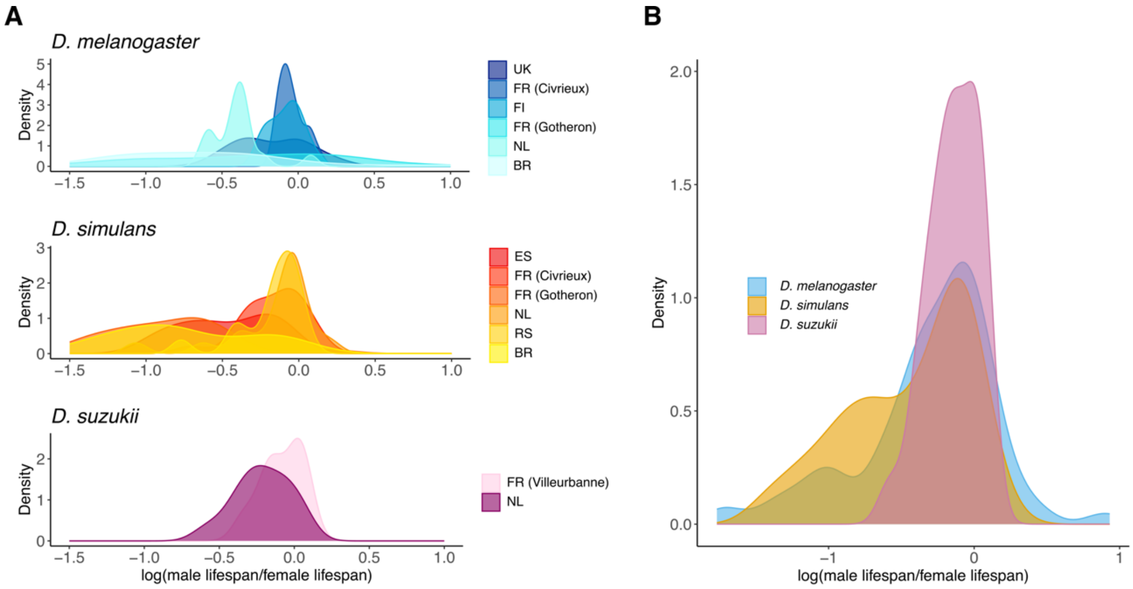

**Figure S2.** PCA analysis plot on normalized counts for genes in **A)** *D. melanogaster*, **B)** *D. simulans*, and **C)** *D. suzukii*. PCA revealed that PC1 accounts for over 60% of the variance and separates samples by sex across all three Drosophila species. PC2 explained 7-15% of the variance, distinguishing samples by age in *D. melanogaster* and *D. suzukii*, and by strain in *D. simulans*.

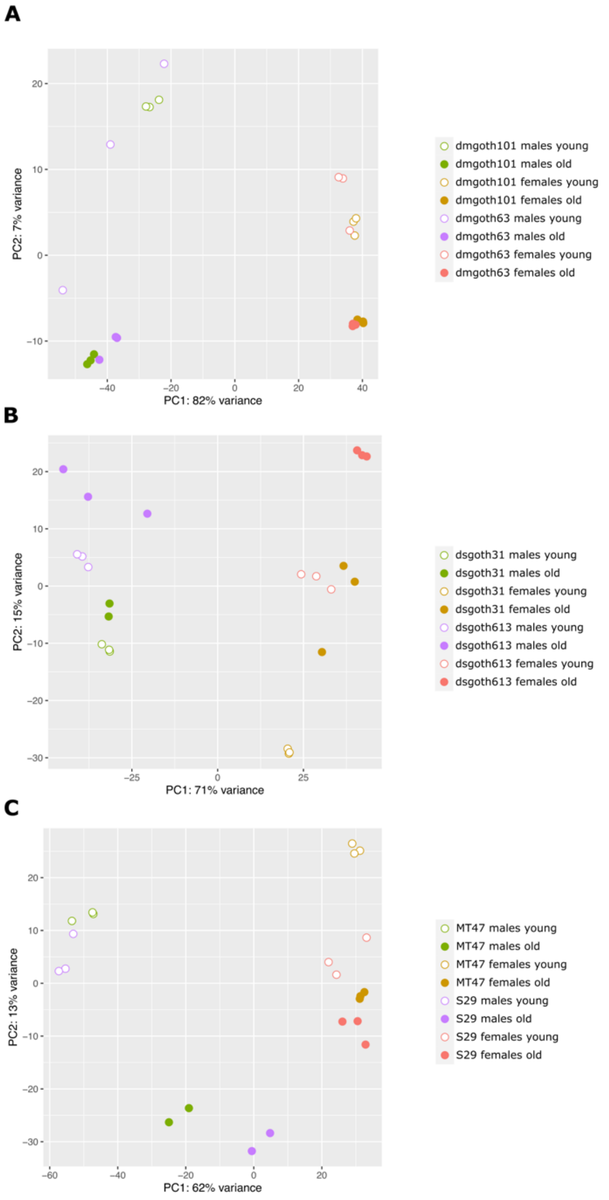

**Figure S3. A)** Volcano plots of DEGs between old and young samples in all the species tested. Statistically significant upregulated genes are depicted in red and statistically significant downregulated genes are in blue. The name of the five most upregulated and downregulated genes in each strain are shown. **B)** Barplots showing the number of upregulated (in red) and downregulated (in blue) genes between old and young samples in all the strains and species tested. Results for males are shown in upper panels while results for females are in lower panels.

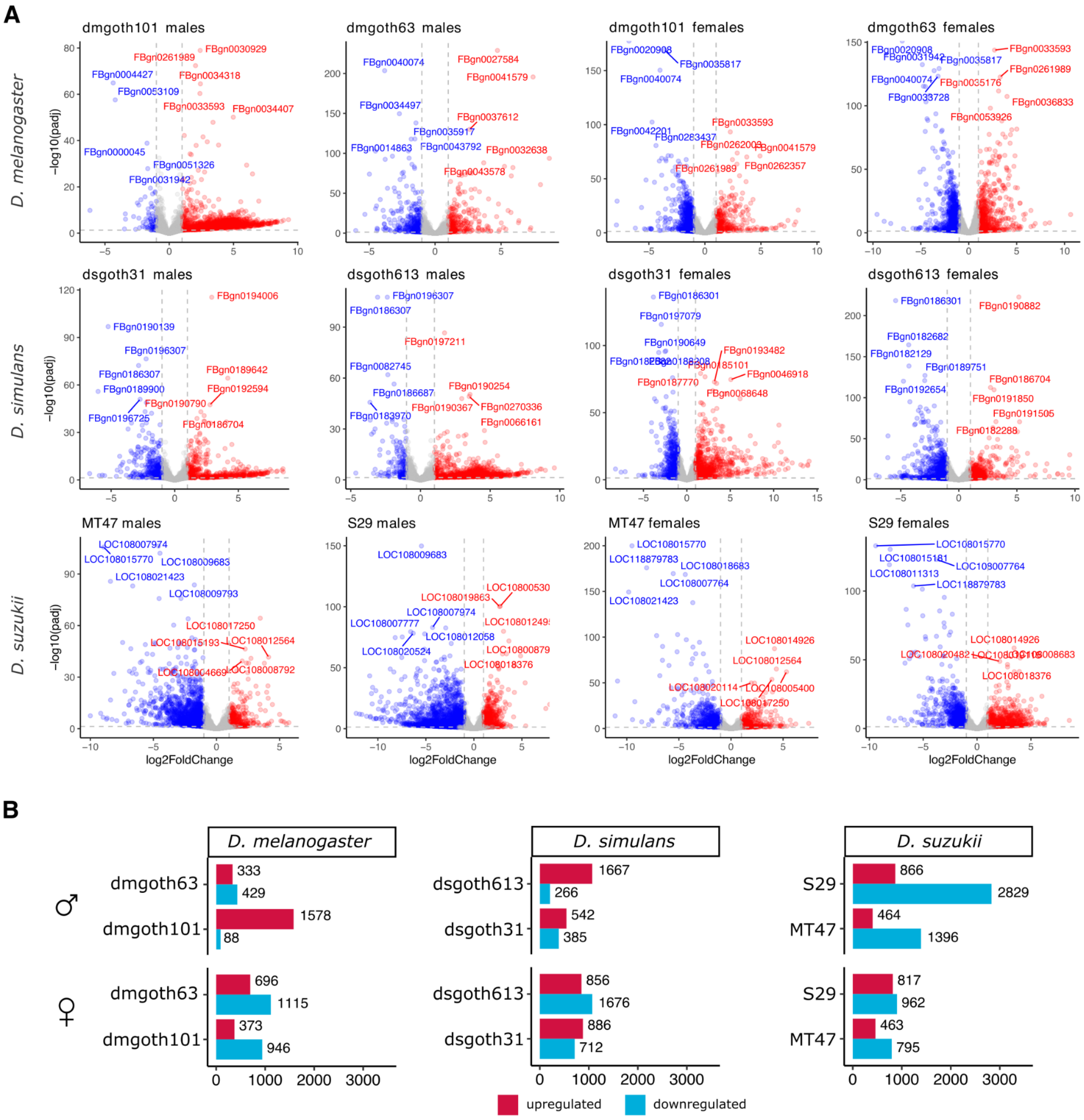

**Figure S4**. Heatmaps showing the expression of previously-associated ageing genes in all the strains from *D. melanogaster*, *D. simulans*, and *D. suzukii*, averaged across replicates. Colors represent log2 fold changes, with red corresponding to upregulated genes and blue corresponding to downregulated genes. Significant p adjusted values are represented with asterisks. A substantial proportion of genes associated with ageing were differentially expressed in the older flies: 14.1%-24.2% in *D. melanogaster* males, 44.3%-45.0% in females; 27.0%-29.6% in *D. simulans* males, 49.3%-53.9% in females; and 25.8%-40.8% in *D. suzukii* males, 25.8%-30.0% in females.

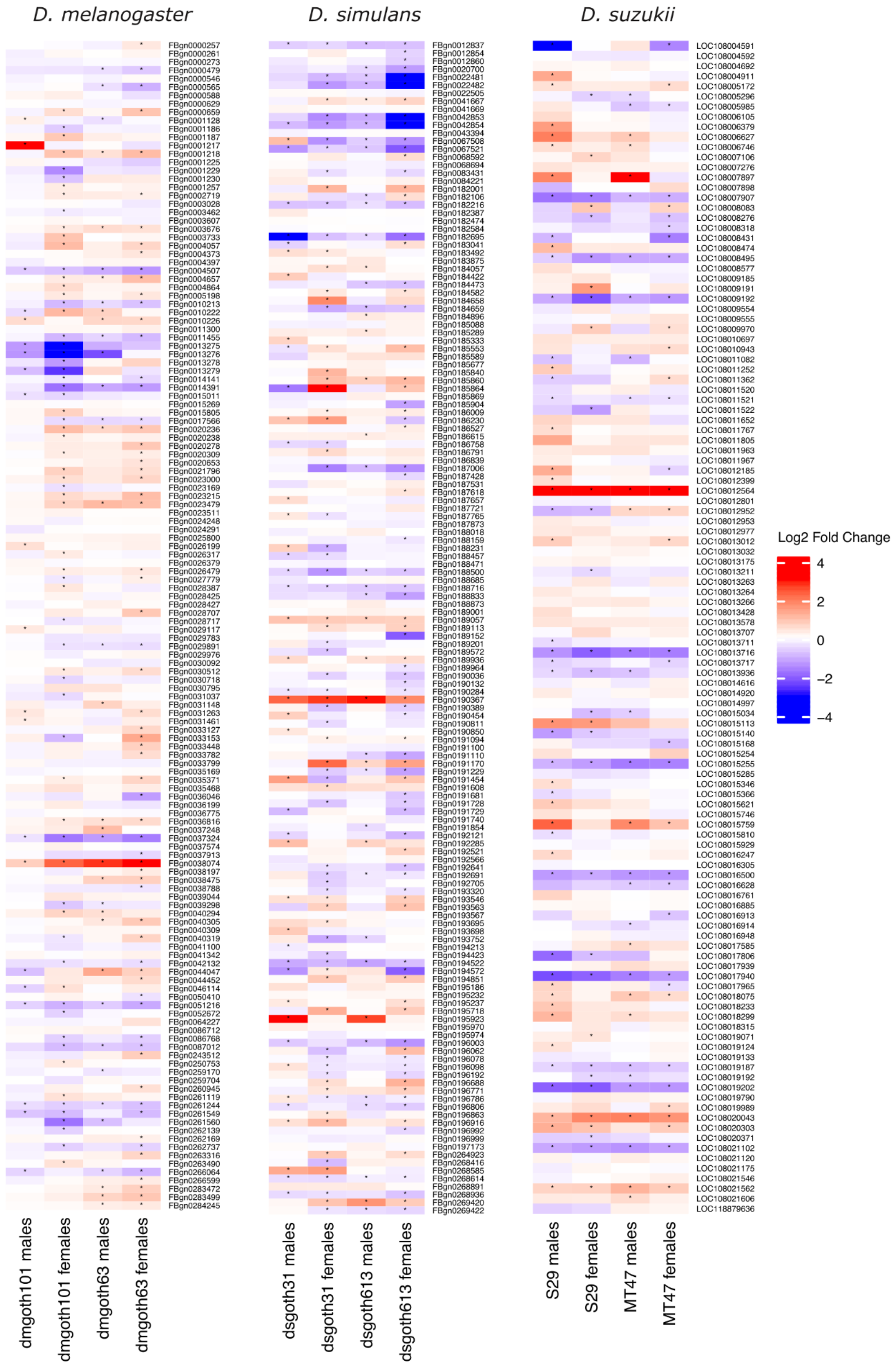

**Figure S5.** Upset plots showing the number of shared DE TE families between samples. Highlighted in red are the number of shared DE TE families between all the samples.

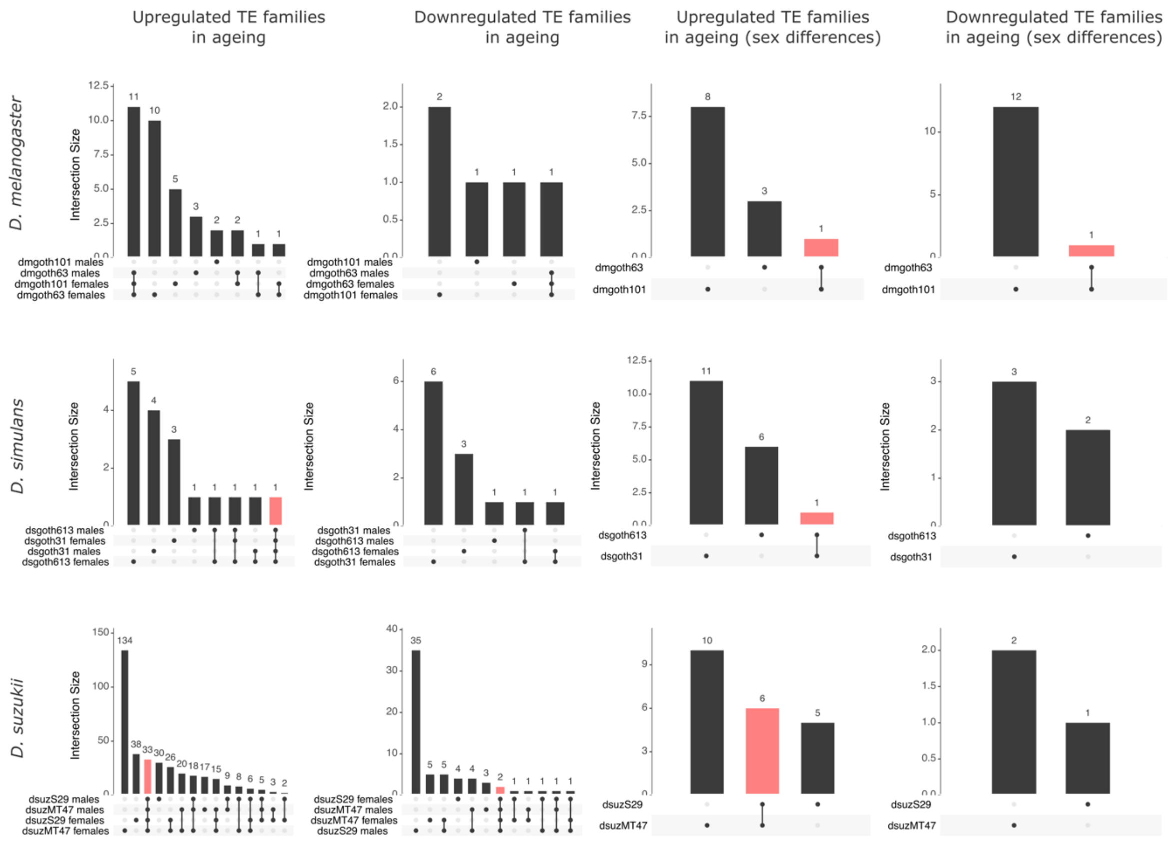

**Figure S6**. Upset plots showing the number of shared DETE copies between males and females from the same strain. Highlighted in red are the number of shared DETE copies between both samples.

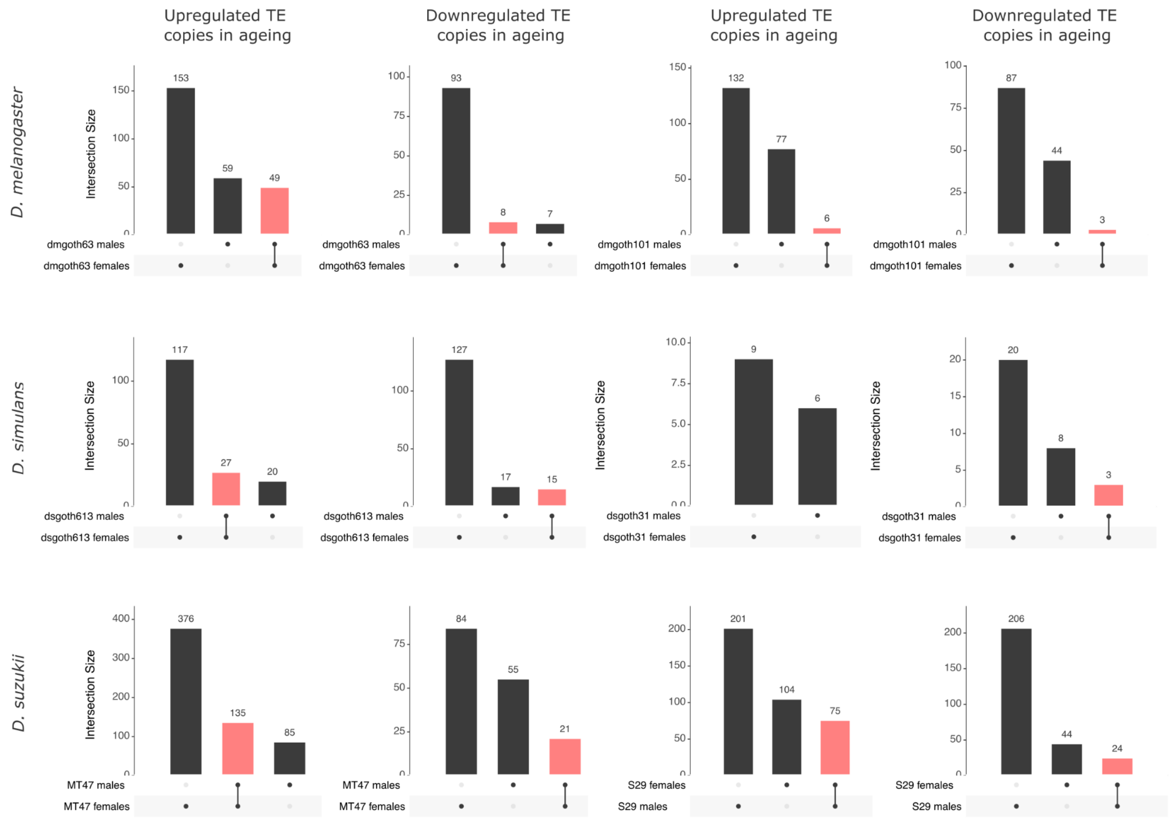

**Figure S7**. Circular track plots showing the distribution of TE insertions (in grey) and upregulated TE copies (in red) in the genome of the *D. melanogaster* strains (plots represented in blue), *D. simulans* (orange), and *D. suzukii* (pink). Plots show the biggest scaffolds assembled for simplification. Circular track plots were generated with the *BioCircos* R package using for each strain its genome assembly and a *bed* file containing the coordinates of the annotated TEs and the upregulated TE copies in old males compared to old females.

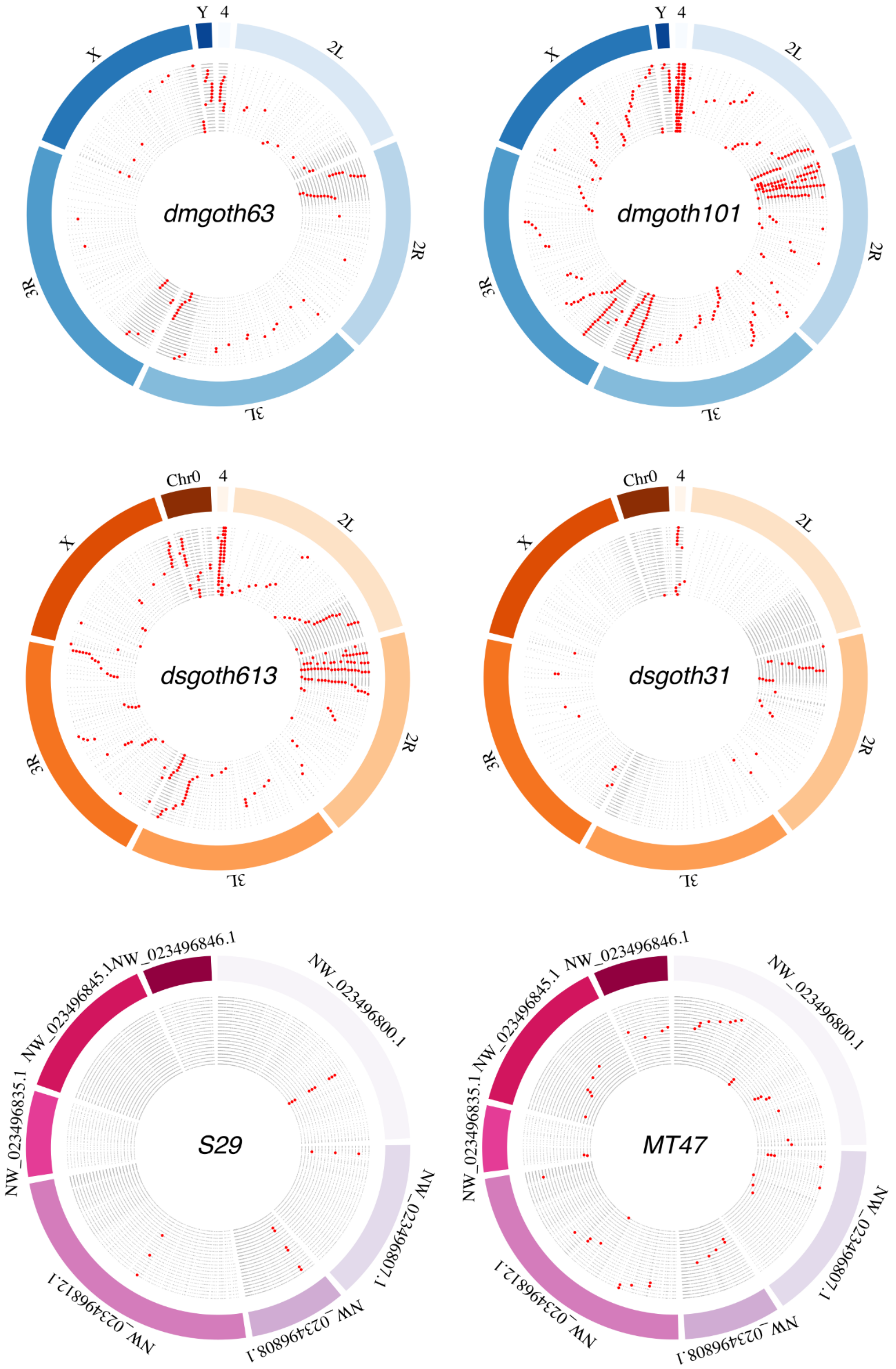

**Figure S8**. Upset plots showing the number of shared chimeric transcripts between sexes and ages from the same strain.

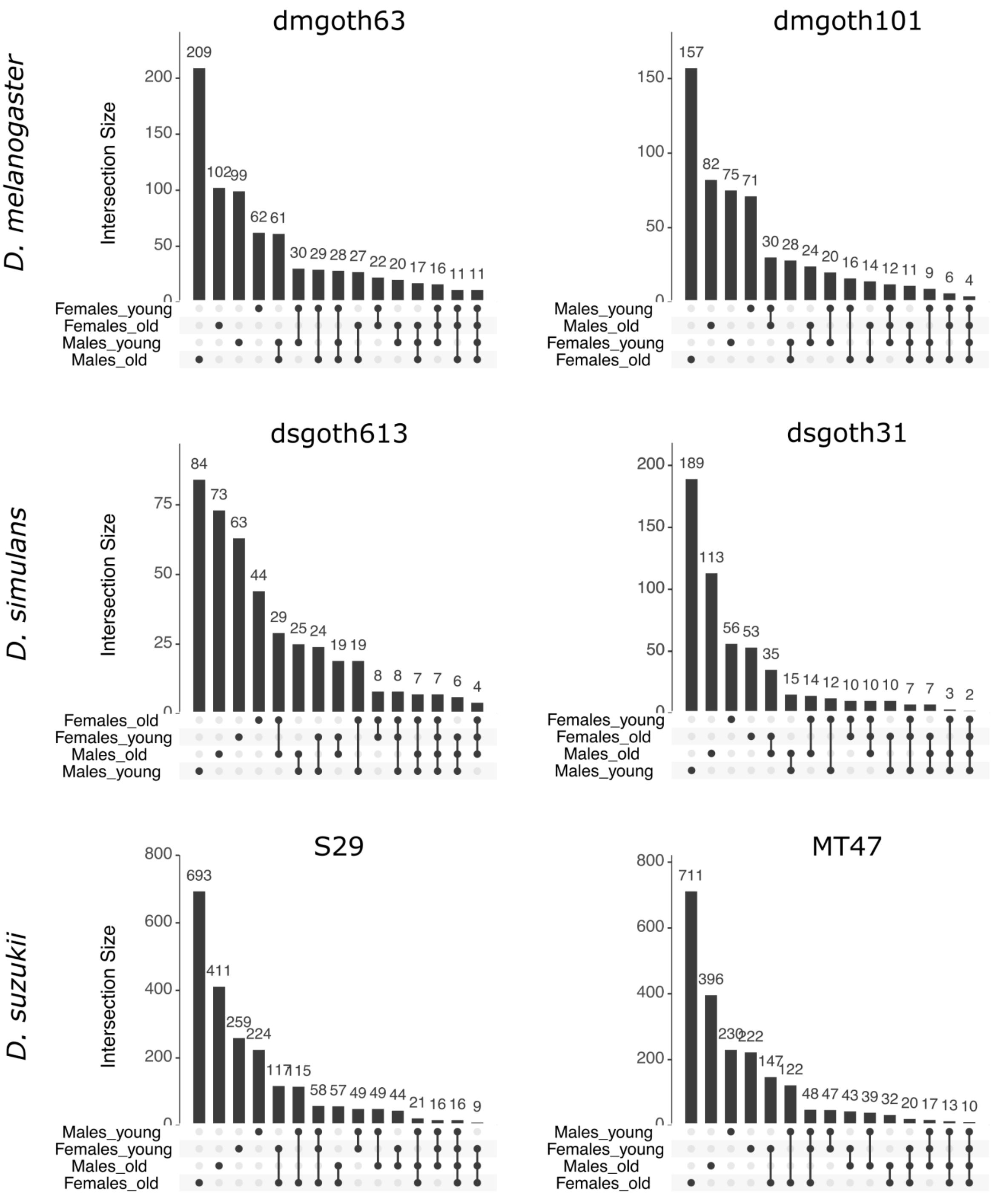

**Figure S9.** Pearson correlations between the TE insertion abundance in the genome for each family and the number of generated chimeric transcripts.

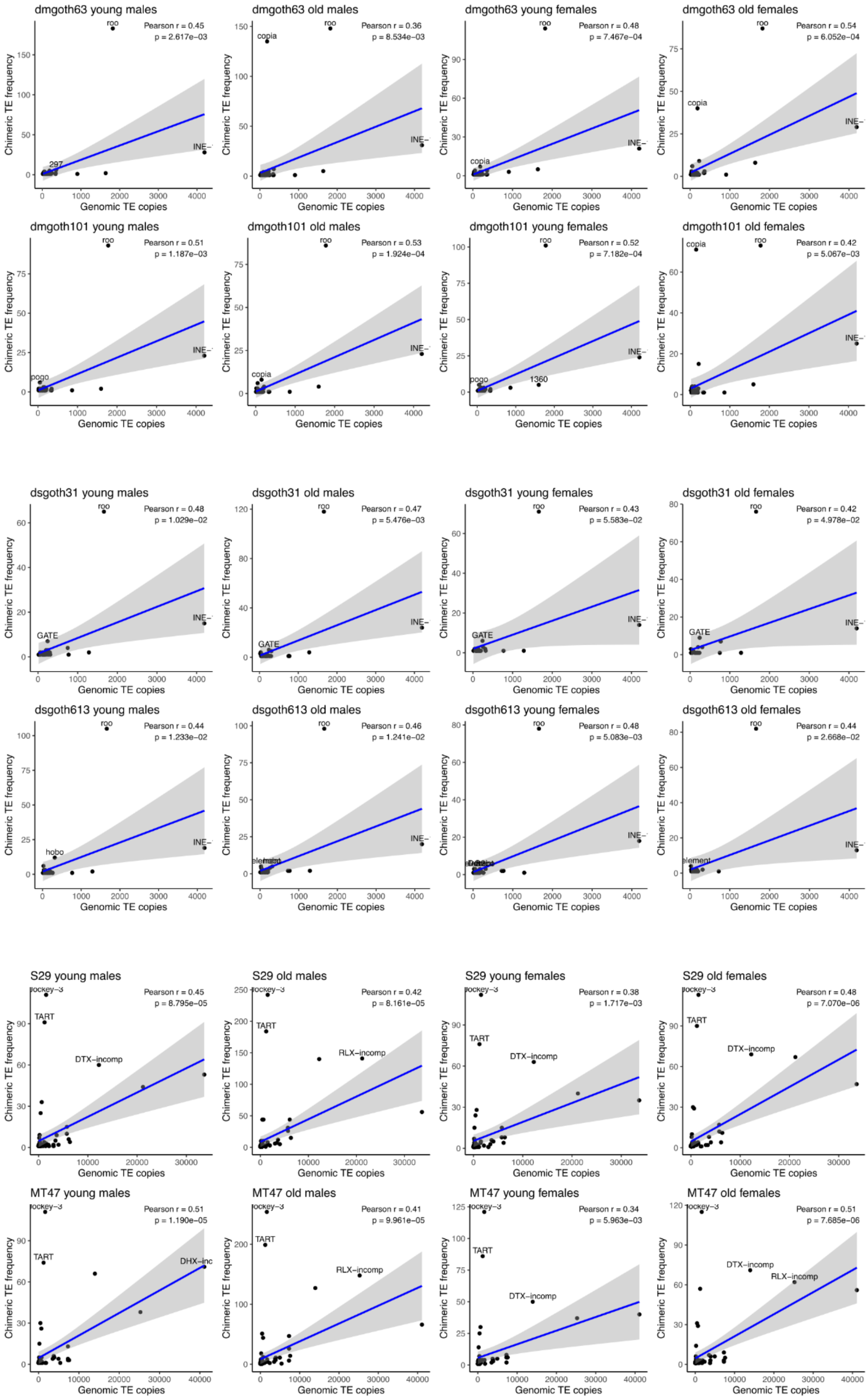

**Figure S10.** Barplots showing the number of chimeric transcripts generated by specific TE families in *D. melanogaster*.

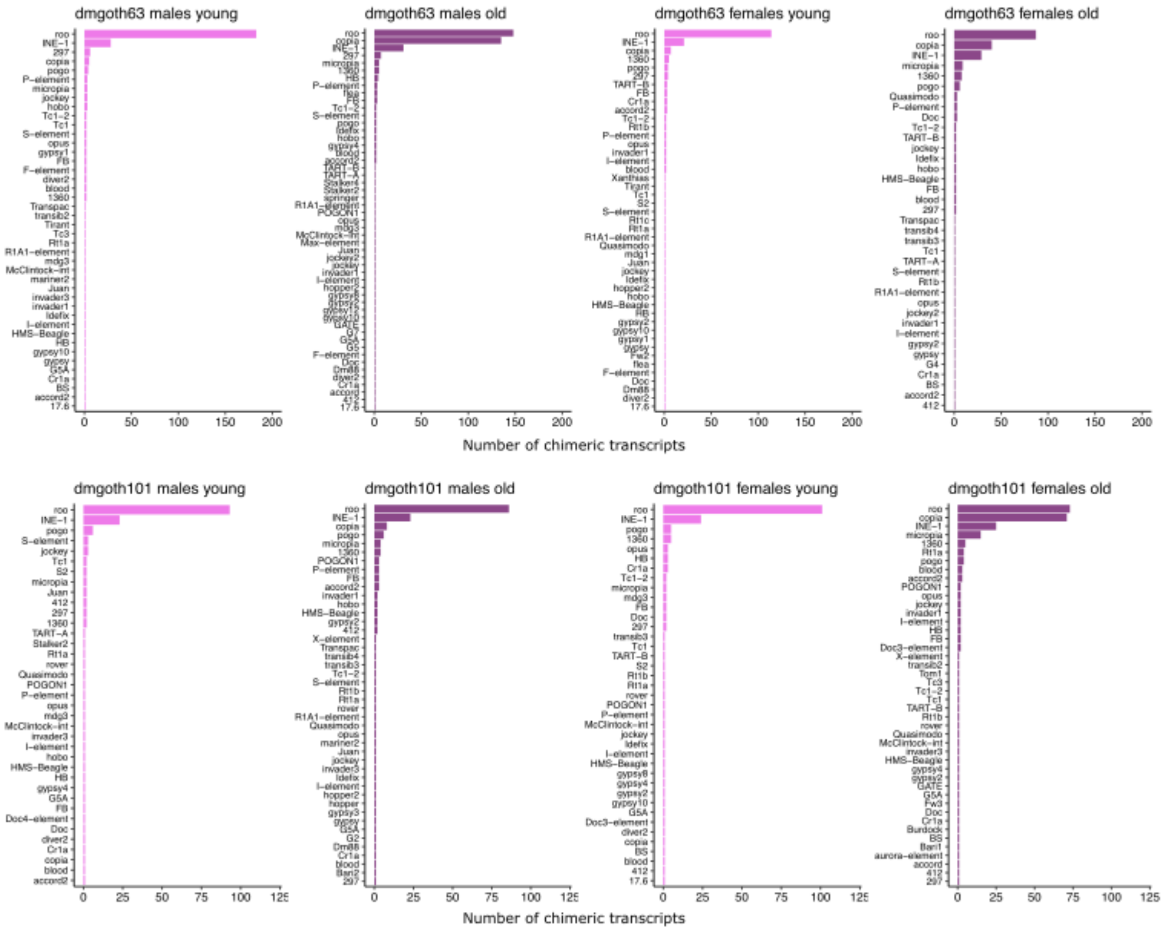

**Figure S11**. Barplots showing the number of chimeric transcripts generated by specific TE families in *D. simulans*.

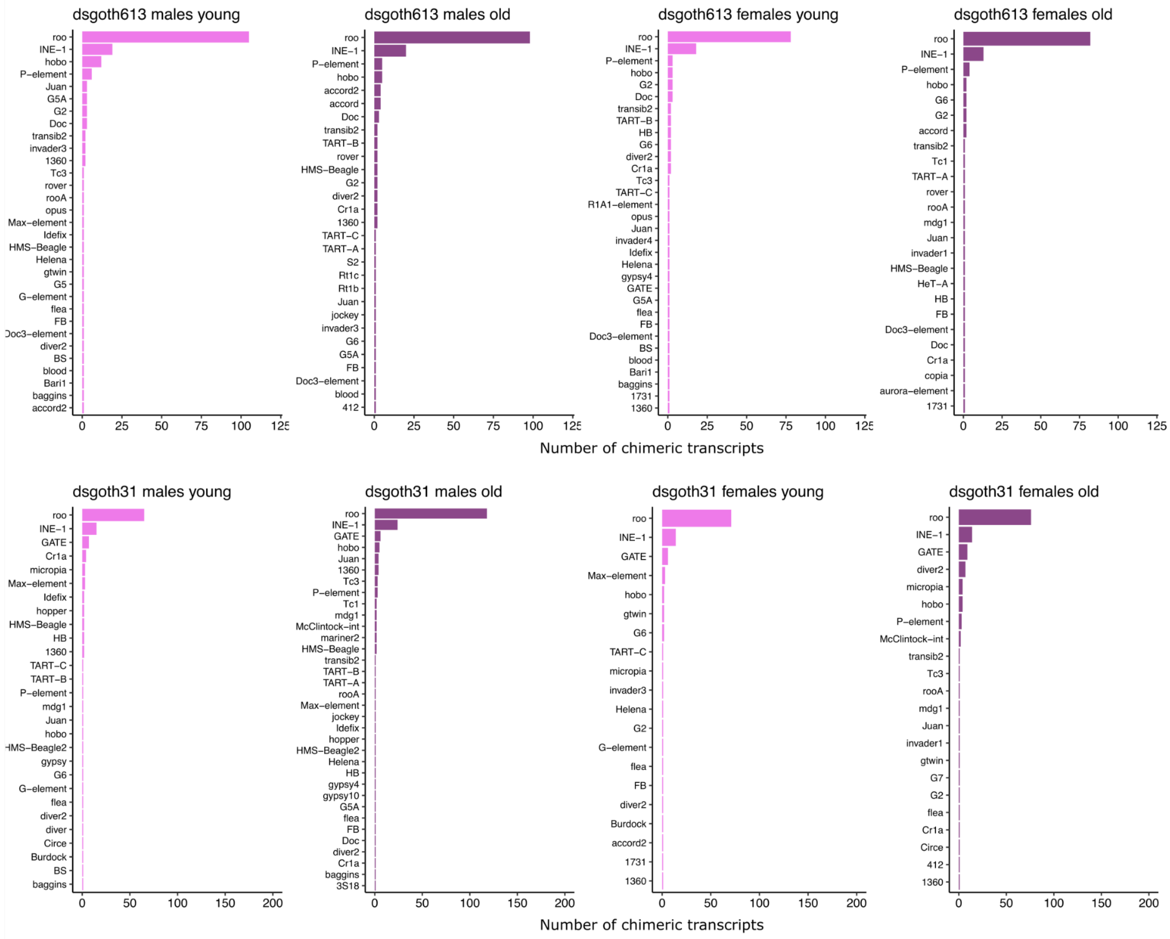

**Figure S12**. Barplots showing the number of chimeric transcripts generated by specific TE families in *D. suzukii*.

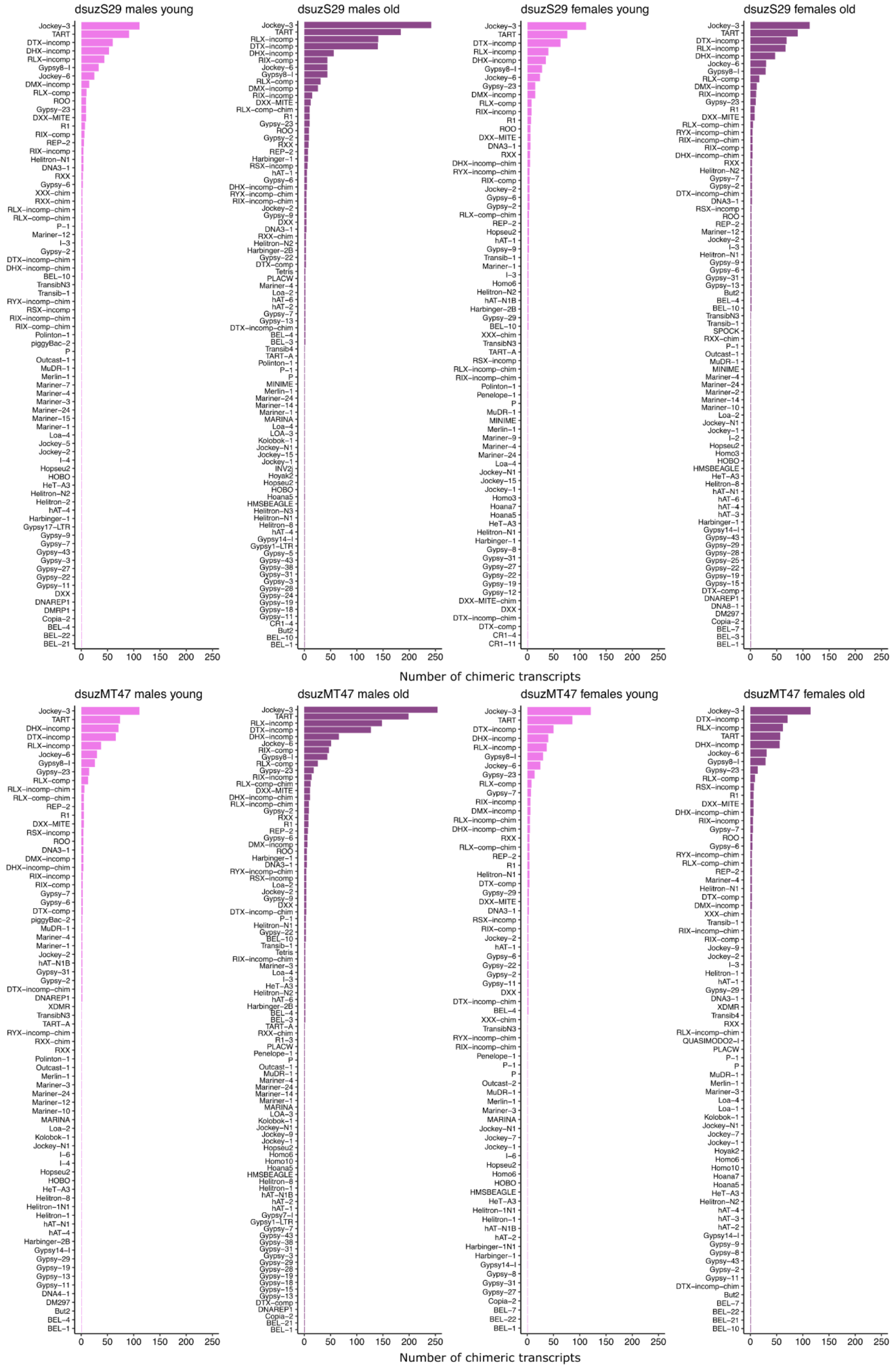

**Figure S13**. **A**) SGL values measured as the logarithm of the ratio between male and female mean lifespan are shown for A reciprocal crosses (left panel) and B reciprocal crosses (right panel) of *D. melanogaster*. Results for maternal strains are shown in grey (dmgoth101) and black (dmgoth63) while different colours represent generations of introgression from F1 to F5. Dots show mean SGL values and lower and upper whiskers indicate the upper and lower 95% confidence intervals. **B**) Kaplan–Meier survivorship curves for males (left panel) and females (right panel) in A reciprocal crosses. **C**) Kaplan–Meier survivorship curves for males (left panel) and females (right panel) in B reciprocal crosses.

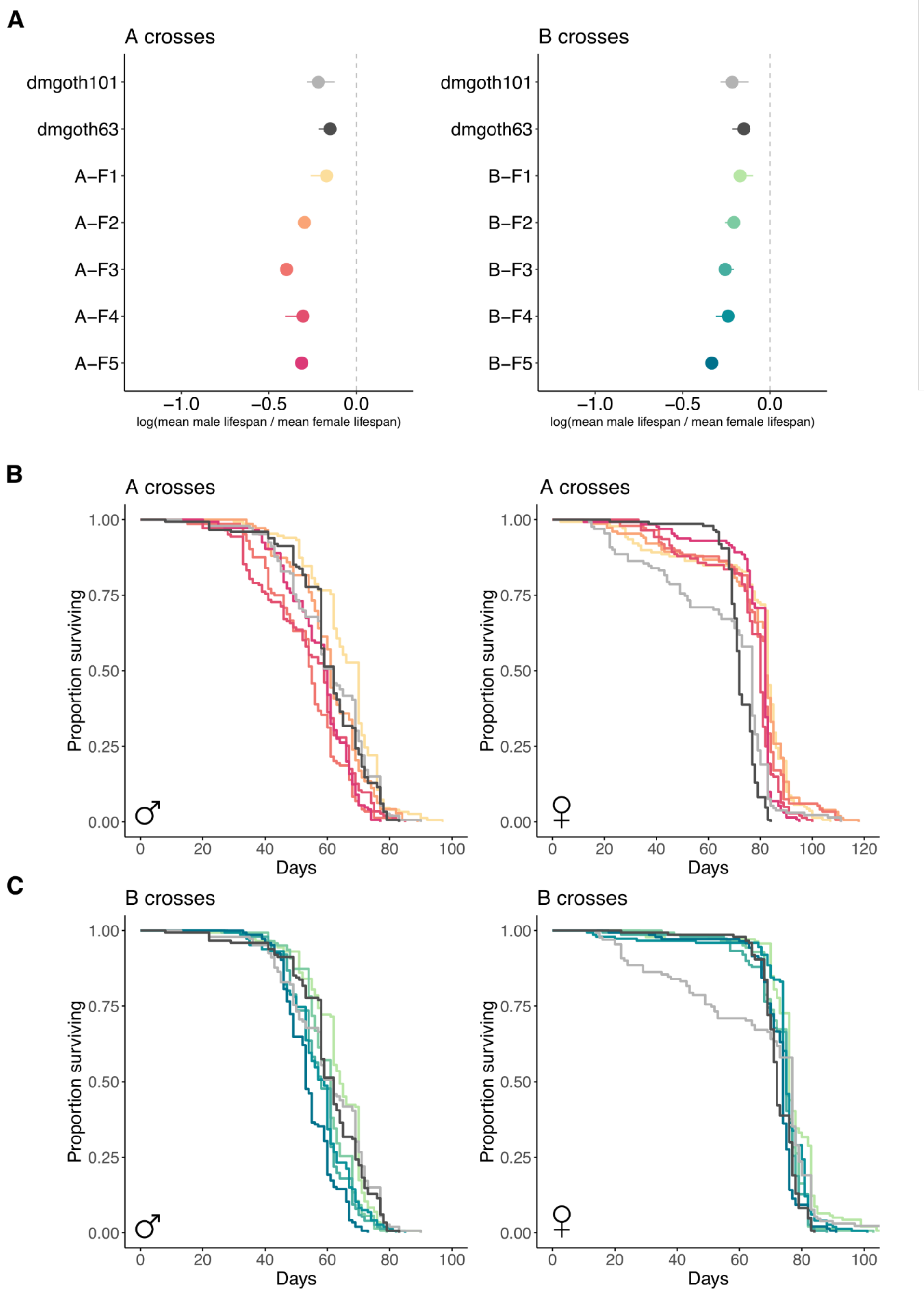
